# The Angiosarcoma Project: enabling genomic and clinical discoveries in a rare cancer through patient-partnered research

**DOI:** 10.1101/741744

**Authors:** Corrie A. Painter, Esha Jain, Brett N. Tomson, Michael Dunphy, Rachel E. Stoddard, Beena S. Thomas, Alyssa L. Damon, Shahrayz Shah, Dewey Kim, Jorge Gómez Tejeda Zañudo, Jason L. Hornick, Yen-Lin Chen, Priscilla Merriam, Chandrajit P. Raut, George D. Demetri, Brian A. Van Tine, Eric S. Lander, Todd R. Golub, Nikhil Wagle

## Abstract

Despite collectively accounting for 25% of tumors in U.S. adults, rare cancers have significant unmet clinical needs as they are difficult to study due to low incidence and geographically dispersed patient populations. We sought to assess whether a patient-partnered research approach using online engagement can overcome these challenges and accelerate scientific discovery in rare cancers, focusing on angiosarcoma (AS), an exceedingly rare sarcoma with a dismal prognosis and an annual U.S. incidence of 300 cases. Here, we describe the development of the Angiosarcoma Project (ASCproject), an initiative enabling patients across the U.S. and Canada to remotely share their clinical information and biospecimens for research. The project generates and publicly releases clinically annotated genomic data on tumor and germline specimens on an ongoing basis. Over 18 months, 338 AS patients registered for the ASCproject, comprising a significant fraction of all patients. Whole exome sequencing of 47 AS tumors revealed several recurrently mutated genes, including *KDR*, *TP53*, and *PIK3CA*. Activating mutations in *PIK3CA* were observed nearly exclusively in primary breast AS, suggesting a therapeutic rationale in these patients. AS of the head, neck, face, and scalp (HNFS) was associated with high tumor mutation burden and a dominant mutational signature of UV light exposure, suggesting that UV damage may be a causative factor in HNFS AS and that this AS subset might be amenable to immune checkpoint inhibitor therapy. Medical record review revealed two patients with HNFS AS received off-label treatment with anti-PD-1 therapy and experienced exceptional responses, highlighting immune checkpoint inhibition as a therapeutic avenue for HNFS AS. This patient-partnered approach has catalyzed an opportunity to discover the etiology and potential therapies for AS patients. Collectively, this proof of concept study demonstrates that empowering patients to directly participate in research can overcome barriers in rare diseases and enable biological and clinical discoveries.

---

Due to low incidence, rare cancer patients are often treated at disparate institutions distributed across the country, ranging from tertiary medical centers to community hospitals. This poses barriers to large-scale scientific studies urgently needed to understand the biology of rare cancers and develop better treatments^1^. We hypothesized that the challenges of rare cancer research could be addressed by engaging patients directly and empowering them to share their samples, their data, and their experiences. In principle, a patient-partnered approach that harnesses the power of social media and patient networks and enables patients to remotely participate irrespective of geography could overcome the barriers of low patient numbers seen at any single institution and aggregate a significant number of rare cancer patients from numerous institutions in a unified clinicogenomic study, thereby rapidly yielding discoveries.

We aimed to test this hypothesis in angiosarcoma, a disease that represents just 1-2% of soft tissue sarcomas, which in turn, comprise less than 1% of adult malignancies^2, 3^. The prognosis for AS is poor, with a reported 5-year disease-specific survival of 38%^2^. Although there have been small genomic studies of AS to date^4–10^, the majority of AS have no known genomic, environmental, or iatrogenic etiology, and effective therapies for most AS patients are lacking.

Working closely with patients and patient advocates, we developed a website (ASCproject.org), which allows AS patients living anywhere in the United States and Canada to register for the ASCproject (Figure 1A). Patients in the angiosarcoma community were deeply involved in all aspects of the project design, implementation, testing, and refinement, including all elements of the study website from images to consent language. AS patients joined the ASCproject rapidly after launch, with 120 patients registering in the first month and a total of 338 patients registering within 18 months (Figure 1B). This represents not only a significant proportion of people living with this disease in the U.S., but also a substantially increased pace of enrollment compared to previous efforts (with the largest previous AS study having collected clinical data from 222 patients treated over 14 years^2^). Online consent for the ASCproject allowed for acquisition of medical records and biological samples (tumor, saliva, and blood), analysis of whole exome sequencing (WES) on tumor and germline DNA, and sharing of de-identified patient-reported, clinical, and genomic data via public databases (Figure 1C-D). Patients continued to be engaged throughout the ASCproject and were regularly provided study updates (Figure 1C).

**Figure 1.**
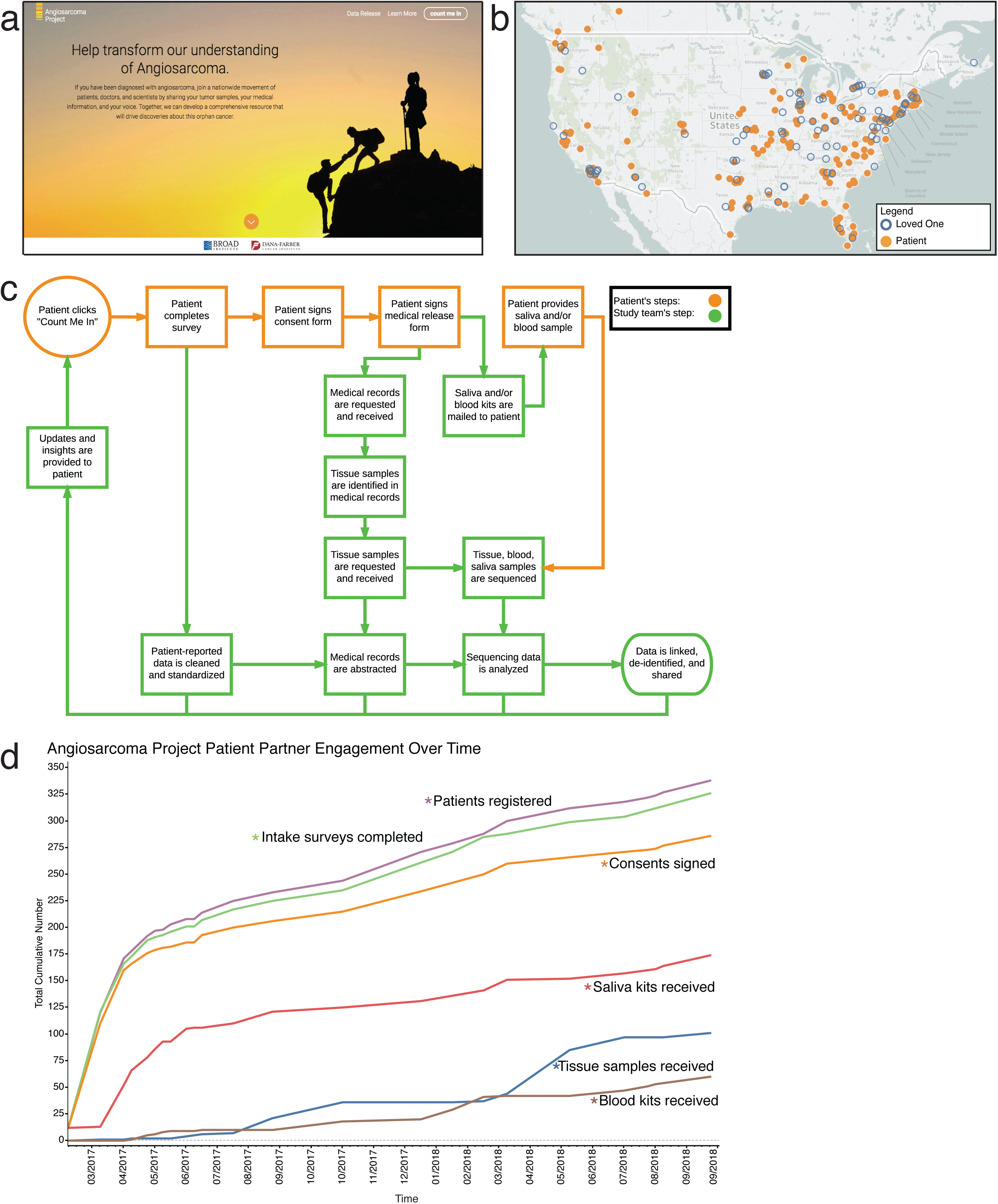
Building a patient-partnered project in Angiosarcoma. (a) Homepage of the ASCproject.org website, which shows images and text designed with AS patients. To begin the project enrollment process, patients click the ‘count me in’ button seen in the upper right corner. (b) Map of the U.S. and Canada showing geographical locations of patients (solid orange circles) and loved ones (open blue circles) who registered for the ASCproject between January 1, 2017 and September 30, 2018. (c) Schematic detailing the process of the Angiosarcoma Project (orange boxes indicate patient steps; green boxes indicate steps taken by study team members). (d) Plot depicting the cumulative totals of patient-driven aspects of the ASCproject between March 2017 and September 2018, which includes numbers of patients registering for the ASCproject (purple), patient intake surveys completed (green), patient consent forms signed (orange), and receipt at the Broad Institute of saliva kits (red), AS tumor tissue samples (blue), and blood kits (brown). 15 AS patients served as beta-testers of the website before the public launch of the ASCproject in March 2017, resulting in non-zero values at March 1, 2017. AS, Angiosarcoma; ASCproject, The Angiosarcoma Project.

Although the study is ongoing, the following analyses were conducted with the 227 patients who had fully consented as of September 30, 2018 (Figure 1D). These 227 patients received care for AS at 340 different clinical institutions, including 289 institutions that were reported only once by any given participant (Supplemental Figure 1) - demonstrating the importance of online platforms to overcome the geographic isolation that has traditionally inhibited large-scale studies in rare cancer patients.

Patients self-reported demographic information, sites of primary AS, as well as other AS and prior cancer information through an intake survey (Figure 2; Supplemental Figures 2 and 3; Supplemental Tables 1 and 2). Patients joining the ASCproject spanned newly diagnosed patients to long-term survivors, with the elapsed time between primary AS diagnosis and ASCproject enrollment ranging from 5 days to 41 years (Figure 2B).

**Figure 2.**
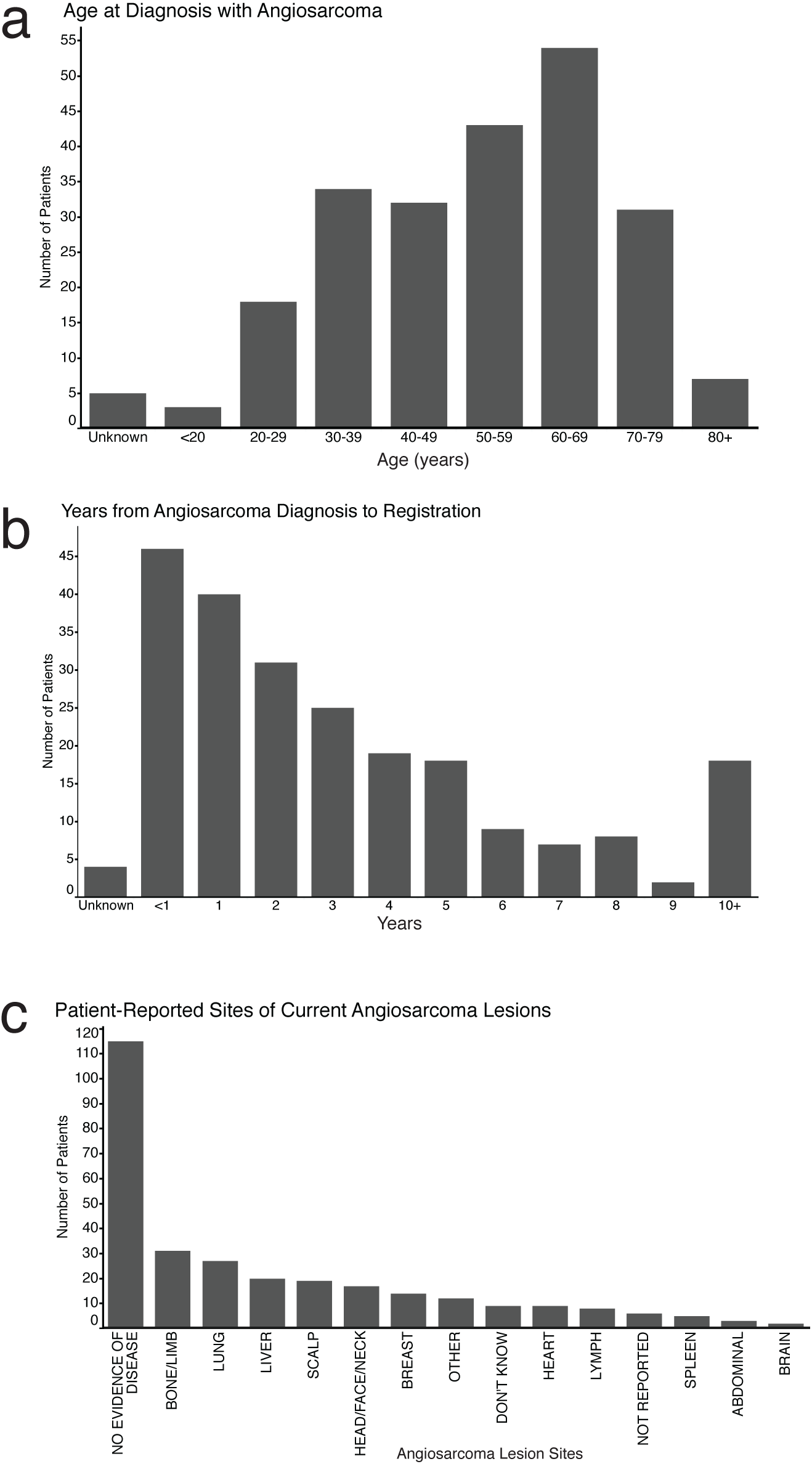
Patient-reported data in the Angiosarcoma Project. Patients first complete an intake survey during the Angiosarcoma Project registration process. Surveys completed by the 227 patients from the U.S. and Canada who consented for the ASCproject as of September 30, 2018 were analyzed. (a) A histogram showing the age in years of patients at initial diagnosis with AS (mean from 222 patients: 53.1 years). These values were calculated from patient-provided date of birth and date of initial AS diagnosis. If insufficient information was provided to calculate this value, patient age was classified as ‘Unknown’ (5 patients). (b) A histogram showing the years elapsed between patients’ initial diagnosis with AS and patients’ registration in the ASCproject (mean from 223 patients: 3.6 years). These values were calculated from the date of project registration and the patient-provided date of initial AS diagnosis. If insufficient information was provided to calculate this value, it was classified as ‘Unknown’ (4 patients). (c) A histogram showing the patient-reported location of angiosarcoma at the time of last intake survey completion. An option was provided for patients to report no evidence of disease. Patients with more than one location of AS were able to provide more than one site. 9 patients responded ‘Don’t Know’, and 6 patients did not respond to this question (‘Not Reported’). ASCproject, The Angiosarcoma Project; AS, Angiosarcoma.

We were able to rapidly acquire medical records and tumor samples from geographically dispersed patients and institutions (Supplemental Figure 4). We performed WES on 70 obtained tumor samples. Forty-seven samples from 36 patients were used for subsequent genomic analysis after assessment of sufficient tumor purity (≥10%) and confirmation as angiosarcoma by centralized pathology review (Supplemental Figure 4). Apart from these considerations, there were no additional selection criteria for these 47 samples. Characteristics of the 36 sequenced patients are shown in Supplemental Figure 5. Abstraction of medical record data (Supplemental Table 3) and histological evaluation were used to classify these tumors into 8 subclassifications of AS (see Supplemental Methods). To our knowledge, this is the largest reported cohort of AS samples that have undergone WES.

We found 30 genes recurrently altered in these 47 samples (determined by somatic alteration frequency, see Methods). This includes genes previously reported as altered in AS^6, 7, 10–12^, as well as several genes that have not been previously reported to be mutated in AS, such as *PIK3CA*, *GRIN2A,* and *NOTCH2* (Figure 3A). Two genes were mutated at a rate significantly higher than expected by chance given background mutational processes (as identified by MutSig2CV^13^; see Supplemental Methods): *TP53* (25%; 9/36 patients) and *KDR* (22%; 8/36 patients) (Supplemental Figure 6). Moreover, mutations in these two genes were also mutually exclusive (p-value=0.02), with 89% (8/9) of *KDR* missense mutations being observed in primary breast AS samples and 82% (9/11) of *TP53* missense mutations detected in AS samples that were not primary breast (Figure 3A; Supplemental Figure 6).

**Figure 3.**
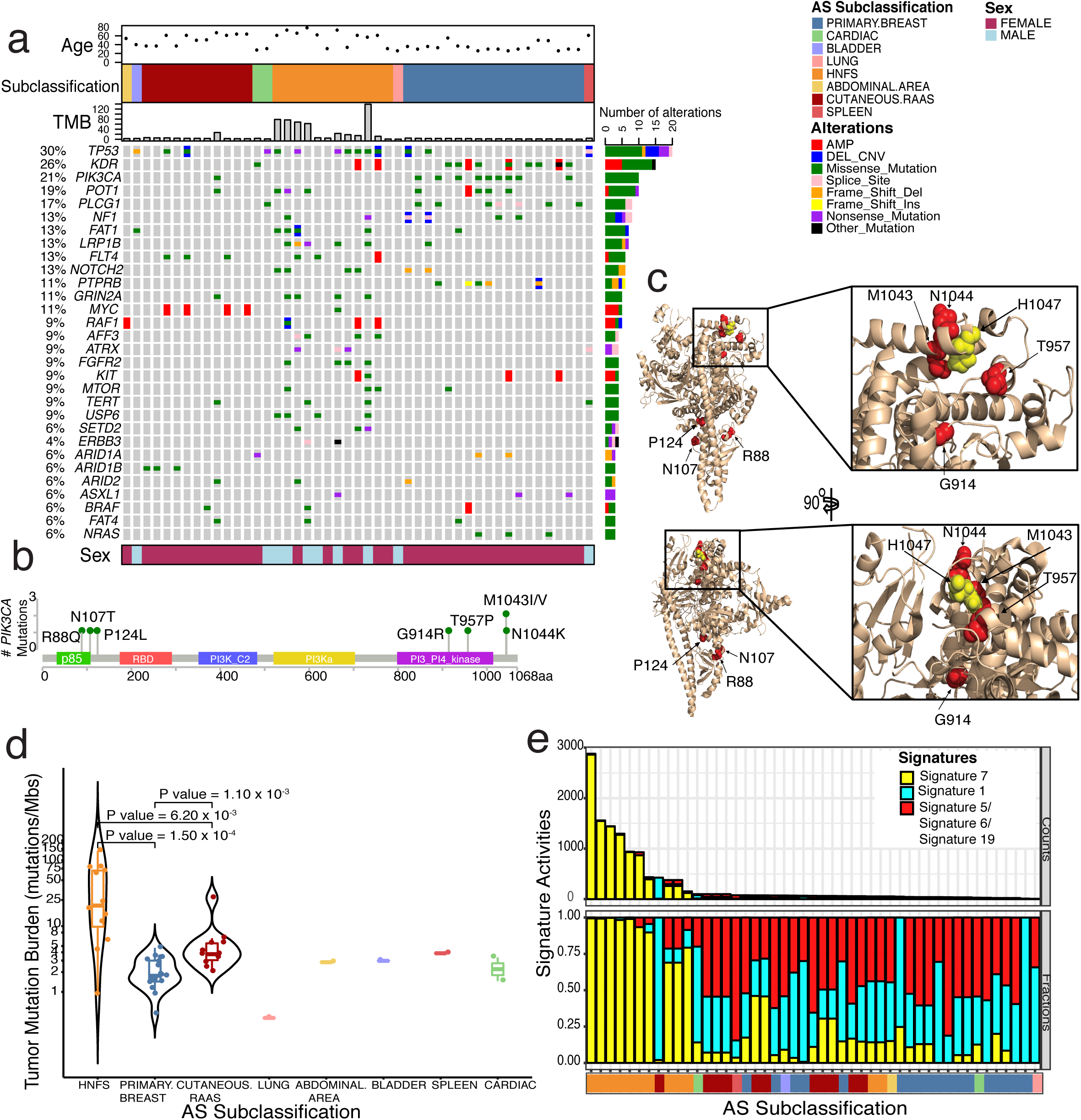
Genomic landscape of angiosarcoma reveals distinct molecular patterns. (a) A co-mutation plot showing somatic alterations and copy number changes in frequently altered genes and other AS associated genes across the cohort of 47 samples from 36 AS patients. Information for each sample is shown in the upper panels including age at AS diagnosis of the patient from whom the sample is derived (top), categorization of each sample to the 8 subclassification of AS (middle), and the tumor mutation burden (TMB) in mutations per megabase (bottom). Patient’s sex is indicated in the lower panel of the plot. (b) Diagram indicating the location and count of mutations occurring in *PIK3CA* in this AS cohort. (c) Crystal structure of p110alpha protein (PDB ID: 3HHM) in wheat cartoon, with red spheres demarcating residues found to be mutated in angiosarcoma tumor samples and the H1047 canonical mutation (yellow spheres). A closer view of the structure with mutations labeled in the regulatory arch region shows a cluster of mutations proximal to H1047 (right hand side box). A 90 degree rotation of this structure is shown in the lower panel. (d) Plot showing the distribution of TMB in mutations per megabase (y-axis; range:0.4-138.9) across tumor samples stratified by the 8 different subclassifications of AS (x-axis). Tumors from the HNFS subclassification exhibit the highest median tumor mutation burden of 20.7 muts/Mbs, which is significantly higher than the median TMB of cutaneous RAAS (3.7 muts/Mbs) and primary breast AS (1.7 muts/Mbs), (P value = 6.20 x 10^-3^ and P value = 1.50 x 10^-4^, respectively). (e) Plot depicting the mutational signature activities across all 47 AS tumor samples. The top panel of counts indicates the total number of mutations (y-axis) attributed to each mutational signature activity within each AS tumor sample (x-axis).The middle panel displays the normalized distribution of signature activities for each sample. The bottom panel shows the associated AS subclassification of each sample. UV light exposure mutational signature (COSMIC Signature 7; indicated in yellow) is dominant in HNFS tumor samples, which also exhibit high tumor mutation counts. AS, Angiosarcoma; HNFS, head, neck, face, scalp; RAAS, radiation-associated angiosarcoma AS; TMB, tumor mutation burden.

*PIK3CA* was one of the most frequently mutated genes in this cohort (21%; 10/47 samples) (Figure 3A). Although alterations in the PI3K pathway have been identified in a previous AS study^14^, mutations in the *PIK3CA* gene itself have not been previously reported in AS to our knowledge. Nine out of the ten *PIK3CA* alterations were found in primary breast AS samples, and this AS subclassification was significantly enriched for *PIK3CA* mutations compared to other subclassifications (9/18 primary breast AS samples versus 1/29 AS samples that were not primary breast; p-value=0.0003) (Figure 3A).

Intriguingly, none of the 8 unique *PIK3CA* mutations we observed were at the canonical *PIK3CA* hotspot residues E545 or H1047^15^ (Figure 3B). Instead, these *PIK3CA* mutations were located in two distinct clusters on the protein structure (Figure 3C), corresponding to regions enriched with activating somatic mutations^16^. Indeed, most of the *PIK3CA* mutations in our AS cohort have been previously described as hotspot mutations in other cancers, and have been shown to be activating *in vitro*^17–19^ (Supplemental Table 4). Moreover, CRISPR experiments in cancer cell lines (depmap.org) demonstrated that lines harboring some of these *PIK3CA* mutations (R88Q, P124L, and G914R) were significantly more dependent on *PIK3CA* than lines with wild-type *PIK3CA* (Supplemental Figure 7). Collectively, these data strongly suggest that the *PIK3CA* mutations detected in this AS cohort are likely to be activating and therefore sensitive to PI3Kα inhibition^20–22^.

*PIK3CA* alterations occur more frequently in breast adenocarcinoma (34.5%) than in other cancer types (>10%)^23, 24^. The fact that different types of activating PI3K mutations are found in breast malignancies with different lineages (angiosarcoma and adenocarcinoma), raises the intriguing possibility that the site of tumor origin (breast), independent of tumor lineage, may be permissive for PI3K pathway activation and aid tumor formation within breast tissue, perhaps due to interaction with the breast microenvironment. Of clinical importance, these observations suggest that PI3Kα inhibitors, one of which was recently approved for the treatment of *PIK3CA*-mutant advanced breast adenocarcinoma^20^, may be useful as a novel targeted therapeutic intervention for these patients with primary breast AS.

We next quantified tumor mutation burden (TMB) for all 47 AS samples. While the median TMB in the full cohort was 3.3 mutations per megabase (muts/Mb), HNFS AS samples showed a significantly higher median TMB than all other AS subclassifications (20.7 muts/Mb for HNFS versus 2.8 muts/Mb for non-HNFS; Wilcoxon’s rank sum test, p-value=0.095; Figure 3D). Moreover, 9 of the 10 samples with high TMB (≥10 muts/Mb) were HNFS AS (Figure 3D). Using mutational signature analysis to understand the possible origins of tumor hypermutation, we found that all 9 of these HNFS samples with high TMB had a dominant mutational signature representing damage from ultraviolet (UV) light (COSMIC Signature 7)^25^ (Figure 3E, Supplemental Figure 6D). The single sample with high TMB that did not have a dominant UV light exposure mutational signature was from a patient with cutaneous radiation-associated AS (C-RAAS) of the breast who also has Lynch syndrome (Figure 3D-E; Supplemental Figure 6D). Our findings suggest that these HNFS AS tumors may have resulted from high TMB caused by UV damage due to sun exposure (Figure 3E, Supplemental Figure 6D). Indeed, 10 out of the 12 HNFS AS tumor samples in this study showed a dominant UV light exposure mutational signature, while none of the other 35 non-HNFS tumor samples harbored this as a dominant mutational signature (p-value=1.27×10^-8^) (Figure 3E, Supplemental Figure 6D). The fact that high TMB and a concomitant dominant mutational signature of UV light exposure occurs uniquely in HNFS AS suggest a common etiologic and genomic basis for HNFS AS, which is an AS subtype accounting for nearly 60% of AS cases^26, 27^.

Since high TMB has been reported as a possible biomarker for response to immune checkpoint inhibition^28–34^, we hypothesized that HNFS AS patients with high TMB might respond particularly well to immune checkpoint inhibitors (ICI). Medical record abstraction of radiation and all systemic treatments for AS received by the sequenced cohort (Figure 4A; Supplemental Figure 8) revealed that 3 of 10 HNFS AS patients had received off-label anti-PD-1 therapy (Figure 4; Supplemental Table 5). Two of those HNFS patients had metastatic AS refractory to standard therapies and demonstrated an exceptional and durable response to pembrolizumab. After receiving several prior therapies for AS that failed, each of these patients has remained disease-free for more than two years after discontinuation of pembrolizumab without receiving any subsequent therapy for AS (Figure 4B, Supplemental Table 6). Of note, these two patients’ tumors had a high TMB (78.5 and 138.9 muts/Mb, respectively; Figure 3D and 4B) and a dominant UV light exposure mutational signature (Figure 3E). The third HNFS AS patient received a single dose of anti-PD-1 treatment, which was stopped due to side effects; this patient went on to take other therapies, none of which resulted in any durable response (Supplemental Table 5).

**Figure 4:**
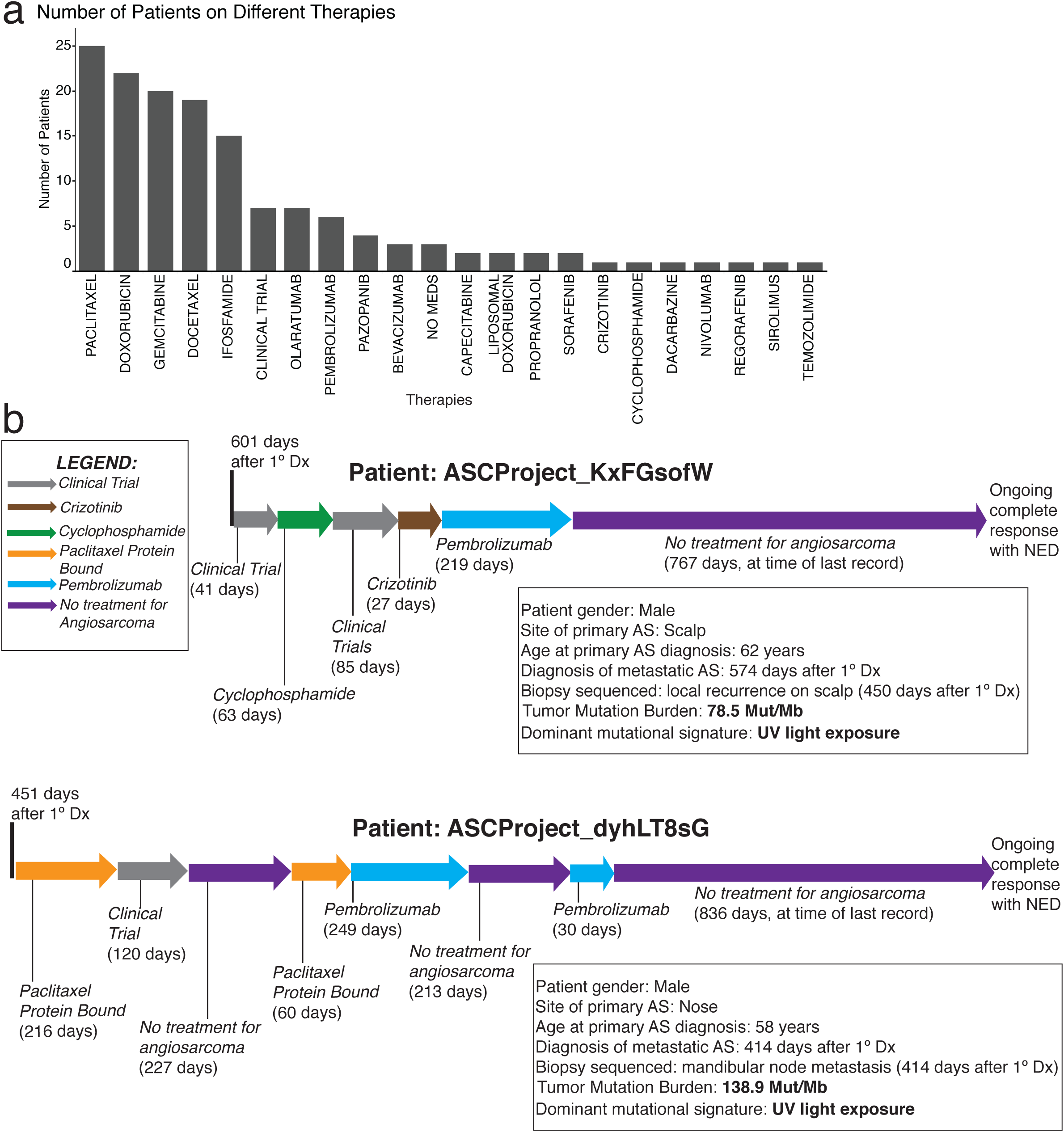
Treatments received by the sequenced AS patient cohort. (a) A histogram showing the number of patients for whom abstracted medical records indicate they received the given treatment for AS listed on the x-axis. This graph depicts the treatments taken by 32 AS patients, as well as 3 patients who received no medications (’No Meds’) per their medical records. There was 1 patient with insufficient medical records to abstract treatment data, who is not included in this chart. (b) Timeline of treatments received by two HNFS AS patients (ASCProject_KxFGsofWAS and ASCProject_dyhLT8sG) in the metastatic setting who each had a complete response to pembrolizumab, as determined by obtained medical record notes. Any time period greater than 200 days in which these patients received no therapy for AS is depicted. These two patients also exhibited high tumor mutational burden and dominant UV light exposure mutational signature. No Meds, No Medications; NED, No Evidence of Disease; AS, Angiosarcoma; 1° Dx, Primary Diagnosis; HNFS, head, neck, face, scalp.

In contrast, 3 of the 26 non-HNFS AS patients (primary breast, cardiac, lung) received off-label use of an anti-PD-1 ICI treatment without clinical benefit (Supplemental Table 5). Tumor samples from these three non-HNFS AS patients had a TMB of less than 5 muts/Mb and did not demonstrate a dominant UV light exposure mutational signature (Supplemental Table 5). These data support the hypothesis that, as in melanoma^35–38^, nearly all HNFS angiosarcoma patients have UV damage-mediated high TMB and might benefit from ICI-directed immunotherapy. Public release of these early results from the ASCproject have helped catalyze the sarcoma community to design clinical trials focused on studying the impact of ICI-immunotherapy in HNFS AS.

In summary, we illustrate that a patient-partnered approach that leverages social media (Supplemental Figure 9) can circumvent the challenges in studying a rare cancer normally encountered through traditional research models and can enable research in the extremely rare cancer angiosarcoma. Within only 18 months of the Angiosarcoma Project launch, we accrued the largest prospective AS cohort that has been reported to date and whose care for AS spanned 340 different institutions. This underscores how this unique research approach can more fully capture and integrate new kinds of valuable data, including off-label use of therapies, that better reflects the varied treatment protocols across different parts of the country, ranging from community hospitals to larger academic medical institutions. The use of online engagement and patient-driven registration may be predicted to skew the demographics of study subjects toward younger^39^ or less sick patients; however, we found that the average age at AS diagnosis in our cohort was just slightly lower than that of a single institution AS study^2^ (53 and 62 years, respectively) and that more than a third of patients (86/227) enrolled within one year of their primary AS diagnosis.

This research approach allowed us to rapidly provide a more detailed clinically annotated-genomic landscape of a rare cancer, AS, which identified significant recurrent genomic alterations. While angiosarcomas have been traditionally classified by site of origin or an environmental or iatrogenic exposure such as prior therapeutic radiation^4–10^, utilizing WES on this sized cohort allowed us to observe additional forms of AS subset stratification that correlate well on a molecular level. Importantly in a malignancy with few effective treatment options, we identified new potential therapeutic strategies for patients with particular AS subclassifications, including primary breast and HNFS AS, which has allowed the sarcoma community to explore developing new clinical trials. To ensure that the ASCproject data can be widely utilized by all researchers, it has been publicly released on cBioPortal.org at regular intervals on a pre-publication basis, with additional data continuing to be released.

The results of the ASCproject suggest that patient-partnered projects may offer a powerful approach for studying cancers. Indeed, the ASCproject is just one of a growing number of patient-partnered projects in different cancers that are part of the *Count Me In* initiative. The ability to rapidly acquire and analyze samples from geographically dispersed patients using the powerful patient-driven approach democratizes research, couples genomic and molecular data to real world patient outcomes, and should be explored in other patient populations that are currently challenging to study through traditional mechanisms.

## Methods

### Website

The ASCproject.org website was developed in collaboration with patients. The website enables angiosarcoma patients across the United States and Canada to learn about the project, register for participation in this research study, sign an electronic informed consent, and provide information about themselves and their disease.

### Informed Consent

Upon completion of online study registration and an intake survey, patients provided informed consent and completed a medical release form in order to be enrolled in the study (Supplemental Table 6). Informed consent was provided by all patients via a web-based consent form as approved by the Dana-Farber/Harvard Cancer Center Institutional Review Board (DF/HCC Protocol 15-057B).

Patient consent allowed the research study team to acquire copies of medical records for abstraction, to send a kit for saliva sample acquisition, to perform sequencing analysis, and to publicly share de-identified linked, clinical, genomic, and patient-reported data. Patients could also opt in to consent to provide a blood sample and/or allow procurement of archived tumor samples for sequencing of germline and tumor DNA.

The analyses conducted for this manuscript were performed with information and samples from patients who consented between January 1, 2017 and September 30, 2018.

### Patient-Reported Data

The patient-reported data for this study consisted of patient responses to the intake survey that accompanied the initial online project registration (Supplemental Tables 1, 7). All 17 questions in this intake survey were optional. Any survey question left blank was categorized as “Not Reported”. Information on format standardization and categorization of patient-reported data is described in the Supplemental Methods.

### Acquisition of Medical Records

For each enrolled patient who completed the medical release form, the study team requested medical records (MR) from all institutions and physician offices from which the patient indicated that they received clinical care. Study staff electronically faxed a detailed MR request form to each facility (Supplemental Table 7). Medical records that had not been received after several months were requested again in the same manner. Medical records were received by fax, mail, or secure electronic message. All medical records were saved to a secure drive.

### Acquisition of Patient Samples

Enrolled patients were mailed separate kits to provide saliva and blood samples.

For saliva samples, patients were asked to provide 2 mL of saliva in the included DNA Genotek (Ottawa, Canada) Oragene Discover (OGR-600) tube^40^ and mail these kits that contain saliva samples in prepaid envelopes to the Broad Institute Genomics Platform (Cambridge, MA).

If the enrolled participant consented to blood samples, patients were sent an empty 10 mL Streck (La Vista, NE) Cell-Free DNA BCT tube^41, 42^ and were asked to bring the tube to their next regularly scheduled clinical appointment and request a courtesy draw. If a courtesy draw was not possible, patients were given the option to go to any Quest Diagnostics^TM^ facility in the U.S. with a voucher for a complimentary blood draw. Kits containing blood samples were mailed back in prepaid envelopes to the Broad Institute Genomics Platform (Cambridge, MA). Blood samples received at the Broad Institute were logged by their unique barcodes and fractionated into plasma and buffy coats. Buffy coats were used to extract germline DNA for WES if no saliva sample was available.

If the participant consented to the acquisition of tumor tissue, portions of stored clinical tumor tissue were requested. A form was faxed to each pathology department requesting one Hematoxylin and Eosin stain (H&E) slide as well as either 5-micron unstained slides (between 8-20 slides) or one Formalin-Fixed Paraffin-Embedded tissue block. Requests explicitly stated that no sample should be exhausted in order to fulfill the request. Tissue samples were received at the Broad Institute by mail. Tissue samples received as blocks were labeled with unique numerical identifiers and cut into three 30-micron scrolls per block, which were then labeled with unique barcode identifiers. Tissue samples received as unstained slides were logged and labeled with unique barcode identifiers. Samples were submitted to the Broad Institute Genomics Platform for sequencing.

### Histological Evaluation

An H&E slide and three additional unstained slides of each tumor sample were sent for centralized expert pathology re-review (JLH) to confirm the diagnosis of angiosarcoma in each sample. Downstream analysis was performed only for samples confirmed to be angiosarcoma.

### Whole Exome Sequencing and Data Analysis

Samples were submitted to the Broad Institute Genomics Platform for processing and sequencing. DNA was extracted from primary and metastatic tumors (for somatic DNA), as well as saliva or blood plasma samples (for germline DNA), and WES was performed, as detailed in the Supplemental Methods. Sequencing data was processed and analyzed to identify somatic single nucleotide variants, small insertions/deletions and copy number alterations using established cancer genomics pipelines at the Broad Institute (see Supplemental Methods). Recurrently altered genes were determined based on frequency of somatic alteration abundance in approximately 680 cancer related genes (https://cancer.sanger.ac.uk/census, https://pathcards.genecards.org/). Mutsig2CV was used to infer significantly recurrent mutated genes in the cohort^13^. Tumor mutation burden (TMB; mutation per megabase) was calculated as the total number of mutations (non-synonymous + synonymous) detected for a given sample divided by the length of the total genomic target region captured with whole exome sequencing^43^. SignatureAnalyzer^44^ was used to identify mutational signatures (as defined by COSMIC) within the cohort, which were further validated using DeconstructSig^45^ for individual tumor samples (see Supplemental Methods). All statistical analysis was performed using R.

### PIK3CA Analysis

To assess *PIK3CA* dependency, CRISPR gene knockout dependency data (Avana dataset), cancer cell line mutation calls, and associated cell line and mutation annotations were taken from the DepMap 19Q1 data release (https://depmap.org/portal/download/). CRISPR knockout gene dependency scores were compared for cell lines with wild-type *PIK3CA* to cell lines harboring hotspot *PIK3CA* mutations and to cell lines with *PIK3CA* mutations observed in this angiosarcoma cohort (see Supplemental Methods). Structural analysis to map *PIK3CA* mutations was performed using PyMOL and the p110alpha protein structure (PDB ID: 3HHM). The GENIE version 5 dataset was used to identify *PIK3CA* mutations across all cancer types (http://genie.cbioportal.org/)^46^.

### Medical Record Abstraction of Clinical Data

Abstraction of each medical record was performed across 40 pre-determined clinical fields independently by two study staff abstractors (see Supplemental Methods). Quality control for concordance was performed by a third abstractor. If required, fields may have received additional review from physicians with expertise in the care of patients with angiosarcoma.

Dates were abstracted to the greatest level of detail available in the record, and all dates reported publicly are based on time elapsed relative to the date of primary diagnosis, as described in Supplemental Methods.

### Data Sharing

The clinically annotated genomic dataset of the Angiosarcoma Project is shared publicly on <u>cBioPortal</u> on an ongoing and regular basis as the data is generated. To protect patient confidentiality, the study dataset is de-identified before it is shared, including the masking of patient IDs and the reclassification of unique patient-reported demographic responses as “other” (see Supplemental Methods).

## Acknowledgements

We thank the many angiosarcoma patients and loved ones of patients who have generously partnered with us to create and drive this research project; we are grateful to work with you every day. We thank the ASCproject advocacy partners (Angiosarcoma Awareness Inc., The Paula Takacs Foundation for Sarcoma Research, SARC, Sarcoma Alliance, Sarcoma Foundation of America, The Sarcoma Coalition, and Target Cancer Foundation). We thank colleagues from across the Broad Institute and Dana Farber for helpful scientific discussions and support. We thank William Hahn for helpful feedback on the manuscript. We thank the Broad Institute Communications & Development teams for their hard work to support this project. We are especially thankful to all members of the *Count Me In* Team, the Wagle lab, the engineering team at the Broad Institute (Andrew Zimmer, Esme Baker, Simone Maiwald, Jen Lapan, Scott Sutherland), the Broad Institute Cancer Program, the Broad Institute Genomics Platform, and the compliance team at the Broad Institute.

