## Supplemental Methods for "The Angiosarcoma Project: enabling genomic and clinical discoveries in a rare cancer through patient-partnered research"

The Angiosarcoma Project is ongoing and continues to enroll patients. The analyses conducted for this manuscript were performed using information and samples from patients who consented between January 1, 2017 and September 30, 2018.

### Project Design:

***Website***

This research project was designed through extensive collaboration with patients in an online angiosarcoma (AS) support group (The Angiosarcoma Project Working Group in Facebook) who generously provided iterative feedback. Working together with patients, a website was developed (ASCproject.org) that enables AS patients across the United States and Canada to learn about the project, register to participate remotely in this research study, sign an electronic informed consent (Supplemental Table 7), and provide information about themselves and their cancer. The Angiosarcoma Project website was publicly launched in March 2017. Prior to that date of the public launch, 15 angiosarcoma patients served as beta-testers of the website. All data collected online as part of the Angiosarcoma Project are stored in a secure database.

***Registration***

Patients registered for the Angiosarcoma Project by providing their first and last name, email address, and confirmation of a diagnosis of angiosarcoma in a primary screening questionnaire. Registered patients were asked to complete a 17-question survey (all questions optional) about their experiences with angiosarcoma (Supplemental Table 7). Registrants acknowledged that their responses will be stored in a secure database. Included in this acknowledgement statement is an understanding that patients may be re-contacted and that they can withdraw from the study at any time by submitting a written electronic request. Withdrawing from the project stops the generation of any new data from a participant's samples, medical records, or additional surveys, but data already generated will not be deleted.

Patients were able to direct questions to study staff at any time (before, during, or after registration or consent) using the contact information (phone number and email) listed on the ASCproject website. Study updates, which included information about the cohort in aggregate, were shared regularly with registered patients through email communications and online social media channels.

This project also has an entry point for family members or loved ones to fill out a survey about the experiences that a patient had before dying from angiosarcoma.

***Consent***

All patients provided written informed consent electronically for this research study as approved by the Dana-Farber/Harvard Cancer Center Institutional Review Board (DF/HCC Protocol 15-057B).

Registrants who submitted the intake survey were asked to electronically sign an informed consent document for formal enrollment in the study (Supplemental Table 7). Email reminders were sent weekly for three weeks to patients who registered but who had not yet completed the consent process. Signing the consent form allowed study staff to obtain medical records and send the patient a saliva kit. Within the consent form, patients also could elect to share a blood sample and a portion of their stored tumor samples. After providing informed consent, patients were then asked to complete a medical release form in order to give their full contact information, as well as a list of physicians and institutions that provided their clinical care for angiosarcoma. Email reminders were sent to consented patients who had not completed the medical release form weekly for three weeks. Upon completion of the consent or medical release forms, signed copies were sent electronically to the enrolled patients for their records.

***Blood and saliva sample acquisition***

Enrolled patients who provided informed consent and also completed the medical release form were mailed saliva kits if the participant reported a valid residential address in the U.S. or Canada. Prior to shipment, saliva kits were labeled with a unique two-dimensional barcode and a pre-paid business reply label addressed to the Broad Institute Genomics Platform (Cambridge, MA). Each barcode identifier was assigned to a unique participant prior to shipment. Participants provided at least 2 mL of saliva in the included DNA Genotek (Ottawa, Canada) Oragene Discover (OGR-600) tube[1] following the included instructions (see link on ascproject.org/data-release) and returned the kits free of charge. Saliva kits received back at the Broad Institute were logged by their unique barcodes and stored at room temperature until advancement to whole exome sequencing (WES).

Blood kits were mailed to registrants who provided informed consent, opted in to this component of the study, and reported a valid residential address in the U.S. or Canada. Blood kits containing an empty 10 mL Streck (La Vista, NE) Cell-Free DNA BCT tube[2,3] were labeled with a unique two-dimensional barcode and included a prepaid FedEx ClinPak envelope addressed to the Broad Institute Genomics Platform (Cambridge, MA) prior to shipment. Each barcode identifier was assigned to a unique participant prior to shipment. Participants provided a blood sample by bringing the kit and instructions (see link on ascproject.org/data- release) with them to their next regularly scheduled clinical appointment and requesting a courtesy draw. If a courtesy draw was not possible, patients were given the option to go to any Quest Diagnostics™ facility in the U.S. with a voucher. This voucher was for a complimentary blood draw for the patient, with the cost covered by the research project. Blood samples received at the Broad Institute were logged by their unique barcodes and fractionated. The resultant plasma and buffy coats were stored at -80$^{\circ c}$. Buffy coats were used to extract germline DNA for WES if no saliva sample was available.

***Medical record and tissue sample acquisition***

Study staff called the offices of the hospitals and physicians listed in each participant's medical release form to confirm the fax number for the medical records department. A detailed request was electronically faxed to each facility that asked for medical records (MR) including clinic notes from treating providers, angiosarcoma treatment data (including radiation and chemotherapy), pathology reports, operative reports, referrals, MD to MD exchange, and genetic testing reports from the date of primary diagnosis through to the date of the request (see Supplemental Table 7 for request form). MR that had not been received after several months were re-requested in the same manner. MR were received by fax, mail, or secure electronic message. All MR were saved to a secure drive to facilitate abstraction. For patients who opted to share portions of their stored tumor samples, the date of each procedure (e.g. biopsy, resection, mastectomy, etc.), the type of procedure, histology, and facility that performed the procedure were abstracted from all pathology reports in order to determine tissue to be requested. Unless the most recent biopsy sample for a given patient measured more than 1 cm, the most recent biopsy samples were not requested in order to avoid exhausting samples that could be needed for future clinical care.

The study staff called the pathology departments associated with each tissue sample to confirm the fax number for tissue requests. A form was faxed to each pathology department requesting 1 Hematoxylin and Eosin stain (H&E) slide as well as either 5-micron unstained slides (a minimum of 8 and a maximum of 20 slides) or 1 Formalin-Fixed Paraffin-Embedded (FFPE) tissue block. Requests explicitly stated that no sample should be exhausted in order to fulfill the request.

Tissue samples were received at the Broad Institute (Cambridge, MA) by mail. Tissue samples received as blocks were labeled with unique numerical identifiers and sent to the Dana-Farber/Harvard Cancer Center Specialized Histopathology Services - Longwood (SHL) Core, which cut three 30-micron scrolls per block, as well as three unstained slides and one H&E for pathology review. SHL was instructed to only cut scrolls if confident that doing so would not exhaust the tissue in the block. The scrolls were then labeled with unique barcode identifiers. Tissue samples received as unstained slides were logged and labeled with unique barcode identifiers.

An H&E slide and three unstained slides from each tumor sample received were sent for expert centralized pathology review to confirm the presence of angiosarcoma in each sample. Genomic analysis and inclusion of a sample in a publicly released genomic dataset occurred only for samples confirmed to be angiosarcoma. If a given tumor sample was not confirmed to be angiosarcoma, that sample was not included in genomic analysis, but the associated patient information was retained and included in the patient-reported dataset of the ASCproject.

Scrolls and unstained slides were submitted to the Broad Institute Genomics Platform for whole exome sequencing. Germline DNA obtained from saliva or buffy coat samples were sent for WES at the same time as its matched tumor sample. Tumor and germline samples that were not matched were not sent for WES.

***Data deposition to public domain***

The clinically annotated genomic dataset of the Angiosarcoma Project is shared publicly in order for all researchers to be able to utilize this data to better understand angiosarcoma. De-identified data has been shared on a recurring basis as the data is generated. Data from the Angiosarcoma Project is currently online at www.cBioPortal.org, and will be shared in additional data repositories as well in the future. For additional details about the Angiosarcoma Project or its data, please contact the study team.

### Patient-Reported Data:

Patient-reported data (PRD) comes from information provided by patients in the 17-question intake survey (Supplemental Table 7), initially completed by patients at the time of project registration. Patients were given the option to update the survey at any time, but were not explicitly asked or required to provide updates after initial survey completion. All survey questions were optional. Any survey question left blank was categorized as "Not Reported" or "No Response". If survey questions included an option to respond by selecting "I don't know" any response of this option was categorized as "Don't Know" ("DK"). If during the completion of the intake survey, any respondent selected "No" to a question that was set up to trigger subsequent survey question(s) that seek more details, the answer to those subsequent questions was marked as "Not Applicable". The Angiosarcoma Project PRD dataset includes information from all consented patients in the United States and Canada, regardless of availability of samples or medical records.

In order to protect confidentiality before any data was shared publicly, race categories that were reported fewer than three times in the dataset were reclassified as "other". When applicable, data elements from PRD were cleaned to standardize formatting and protect patient confidentiality. For questions in which responses indicated a location in the body, 11 anatomical locations were provided as answer choices (Supplemental Table 7), as well as an option to select "Other" and provide a free text response. Any free text responses provided by patients that did indicate a location in the body from 1 of the 11 provided anatomical locations were classified into that category as part of the cleaning process. Those answers that did not fall into one of the categories or had no subsequent free text were classified as "Other". Patient responses to their referral sources to the ASCproject (Supplemental Figure 9) were manually reviewed and assigned into the following categories "Social Media", "Internet Search/Website", "Doctor", "Friend/Family", "Advocacy Partner", and "Other". Additional information regarding cleaning of PRD is available upon request.

**Medical Record Abstraction:**

For each consented patient living in the U.S. or Canada, medical records were requested from all doctors and institutions based in the U.S. and Canada that were listed by the patient in their medical release form. Records were scanned and securely stored by study staff upon receipt. Using medical records, 40 pre-determined clinical fields were manually abstracted for each patient. These clinical fields are listed in detail in a data dictionary that was developed with the input from experts in the field of angiosarcoma (data dictionary available upon request). To help facilitate manual abstraction, scanned medical records were converted to searchable PDF files by using the Optical Character Recognition (OCR) engine known as Tesseract (LSTM model inside Tesseract version 4.0; (https://github.com/tesseract-ocr/tesseract)).

Each medical record was fully abstracted independently by two different abstractors on the study staff. Quality control (QC) for concordance was performed by a third abstractor. During the QC process, the third abstractor compared all abstracted clinical fields from the two other abstractors. For any clinical fields lacking 100% complete concordance, the QC abstractor went back to the record to review. If the QC abstractor was unable to determine the correct answer, these fields received additional review from physicians with expertise in the care of patients with angiosarcoma.

Dates were abstracted to the greatest level of detail available in the record. Dates reported in the medical record only as a year were abstracted as the first of the year. Dates reported in the medical record only as a month and year were abstracted as the first of the month. Dates that could be inferred based on notes were indicated as estimated and not directly abstracted. In order to protect patient confidentiality, all dates reported were based on elapsed time relative to the date of primary diagnosis. Ages were grouped into bins for public data releases.

**Subclassifications of angiosarcoma:**

For patients whose tumor samples underwent sequencing, their first angiosarcoma diagnosis was categorized into eight AS subclassifications using information abstracted from medical records. These eight subclassifications of AS were defined based on guidance provided by angiosarcoma experts (abdominal area, bladder, cardiac, cutaneous radiation-associated, HNFS (head, neck, face, scalp), lung, primary breast, and spleen).

To assign the cutaneous radiation-associated AS (C-RAAS) subclassification, patients' medical records needed to explicitly indicate that radiation was previously administered (during the treatment of a prior cancer) to the proximal area later affected by angiosarcoma. Additionally for the C-RAAS subclassification, these patients' pathology reports were used to confirm cutaneous AS. For the HNFS AS subclassification, all of the patients' records were reviewed and this reconfirmed that all had cutaneous lesions. The primary breast subclassification included AS patients with breast AS that did not receive prior radiation therapy to the proximal area of the breast.

**Biological Sample Processing and Sequencing:**

***DNA Isolation***

DNA was extracted via the Chemagic MSM I with the Chemagic DNA Blood Kit-96 from Perkin Elmer. This kit combines a chemical and mechanical lysis with magnetic bead-based purification. Saliva samples were incubated at 50°C for 2 hours. The saliva was then transferred to a deep well plate placed on the Chemagic MSM I. Whole blood samples were incubated at 37°C for 5-10 minutes to thaw. The blood was transferred to a deep well plate with protease and placed on the Chemagic MSM I. The following steps were automated on the MSM I. M-PVA Magnetic Beads were added to the saliva or the blood and protease solution. Lysis buffer was added to the solution and mixed. The bead-bound DNA was then removed from solution via a 96-rod magnetic head and washed in three Ethanol-based wash buffers to eliminate cell debris and protein residue. The beads were then washed in a final water wash buffer. Finally, the beads were dipped in elution buffer to resuspend the DNA sample in solution. The beads were then removed from the solution, leaving purified DNA eluate. DNA samples were quantified using a fluorescence-based PicoGreen assay. For FFPE tumor tissues, DNA and RNA were extracted simultaneously using Qiagen's AllPrep DNA/RNA FFPE kit.

***Library Construction***

Library construction was performed as described[4], with the following modifications: initial genomic DNA input into shearing was reduced from 3$\mu$g to 10-100ng in 50$\mu$L of solution. For adapter ligation, Illumina paired end adapters were replaced with palindromic forked adapters, purchased from Integrated DNA Technologies, with unique dual-indexed molecular barcode sequences to facilitate downstream pooling. Kapa HyperPrep reagents in 96-reaction kit format were used for end repair/A-tailing, adapter ligation, and library enrichment PCR. In addition, during the post-enrichment SPRI cleanup, elution volume was reduced to 30$\mu$L to maximize library concentration, and a vortexing step was added to maximize the amount of template eluted.

***In-solution hybrid selection***

After library construction, hybridization and capture were performed using the relevant components of Illumina's TruSeq Rapid Exome Kit and following the manufacturer's suggested protocol, with the following exceptions: first, all libraries within a library construction plate were pooled prior to hybridization. Second, the Midi plate from Illumina's TruSeq Rapid Exome Kit was replaced with a skirted PCR plate to facilitate automation. All hybridization and capture steps were automated on the Agilent Bravo liquid handling system.

***Preparation of libraries for cluster amplification and sequencing***

After post-capture enrichment, library pools were quantified using qPCR (automated assay on the Agilent Bravo), using a kit purchased from KAPA Biosystems with probes specific to the ends of the adapters. Based on qPCR quantification, libraries were normalized to 2nM, then denatured using 0.2N NaOH on the Hamilton Starlet. After denaturation, libraries were diluted to 20pM using hybridization buffer purchased from Illumina.

***Cluster amplification and sequencing***

Cluster amplification of denatured templates was performed according to the manufacturer's protocol (Illumina) using HiSeq 4000 cluster chemistry and HiSeq 4000 flowcells. Flowcells were sequenced on v1 Sequencing-by-Synthesis chemistry for HiSeq 4000 flowcells. The flowcells were then analyzed using RTA v.1.18.64 or later. Each pool of whole exome libraries was run on paired 76bp runs, reading the dual-indexed sequences to identify molecular indices and sequenced across the number of lanes needed to meet coverage for all libraries in the pool.

### Sequencing Data Analysis:

Please contact for additional details regarding pipeline information.

***Sequence data processing and quality control***

Whole exome sequences were captured using Illumina technology and the sequence data processing and analysis was performed using the Picard and Firehose pipelines at the Broad Institute. The Picard pipeline (http://picard.sourceforge.net) was used to produce a BAM file with aligned reads. This includes alignment to the GRCh37 human reference sequence using the BWA aligner[5] and estimation and recalibration of base quality score with the Genome Analysis Toolkit (GATK)[6]. All sample pairs passed through the Firehose pipeline were subjected to QC testing to test for any tumor/normal and inter-individual contamination as previously described.[7,8]

***Somatic alterations assessment***

The MuTect algorithm was used to identify somatic mutations[8]. To reduce false positive calls, additional analysis of reads covering sites of a putative somatic mutation was done and were realigned with NovoAlign ([www.novocraft.com](http://www.novocraft.com)). An additional iteration of MuTect inference on newly aligned BAM files was performed. Furthermore, false-positive somatic mutation calls were filtered using a panel of normals (PoN), generated from the TCGA dataset and oxoG filter[9]. Small somatic insertions and deletions were detected using Strelka algorithm[10], MuTect 2 and SvABA (https://github.com/walaj/svaba). Insertions and deletions which were called by two out of the three methods mentioned above were used for analysis. Somatic mutations including single-nucleotide variants, insertions, and deletions were annotated using Oncotator[11]. The germline somatic variants were analyzed using the HaplotypeCaller module of GATK[6].

***Mutation Significance Analysis***

Significance of identified somatic mutations was analyzed using MutSig2CV[12], which uses patient and gene- specific mutation rates to estimate a background model of predicted mutation incidence across the genome. MutSig2CV then factors in biological co-variates such as replication timing and gene-expression level on a gene-by-gene basis to account for the increased mutational rate of certain classes of genes.

***Copy number alterations analysis***

ReCapseg was used to analyze somatic copy number alterations (SCNA) from whole exome data. ReCapseg assesses homolog-specific copy ratios from segmental estimates of multipoint allelic copy ratios at heterozygous loci incorporating the statistical phasing software (BEAGLE) and population haplotype panels (HAPMAP3)[13,14]. For copy number alteration significance analysis, segmented copy number data was analyzed by GISTIC 2.0, to identify significantly recurring focal and arm-level amplification/deletion peaks[15].

***Purity and ploidy estimation***

Allele-specific SCNAs and tumor ploidy/purity status were assessed using ABSOLUTE[16]. Angiosarcoma tumor samples which had a purity of 10% or more were submitted to cBioPortal (http://www.cbioportal.org/index.do) and were used for downstream analysis.

***Assessment of Tumor Mutation Burden (TMB)***

Tumor mutation burden (mutation per megabase) was calculated as the total number of mutations (non-synonymous + synonymous) detected for a given sample divided by the length of the total genomic target region captured with the whole exome sequencing[17]. Samples with TMB ≥ 10 mutations per megabase were classified as hypermutated[18].

***Mutational Signature Analysis***

Mutational signatures were analyzed using Signature Analyzer tool[19]. Contributions of different mutation signatures were identified for each sample according to distribution of the 6 substitution classes and the adjacent bases immediately 5′ and 3′ of the mutated base, producing 96 possible mutation subtypes. The extracted signatures were compared against the known and validated signatures[20]. A sample was determined to have a dominant signature based on the maximum signature score attributable to that sample. DeconstructSig[21] was used for additional validation of these identified signatures on an individual tumor sample level.

### ***PIK3CA* Analysis:**

#### ***PIK3CA Structural Analysis***

Structural analysis was performed using PyMOL (The PyMOL Molecular Graphics System, Version 1.2r3pre, Schrödinger, LLC.). The existing structure of the p110alpha protein (PDB ID: 3HHM) was used to map the mutations in *PIK3CA* that were observed in breast angiosarcoma samples.

#### ***Dependency data analysis for PIK3CA mutations***

CRISPR knockout (KO) dependency data (Avana dataset), cancer cell line mutation calls, and associated cell line and mutation annotations were taken from the DepMap 19Q1 data release (https://depmap.org/portal/download/). The Avana dataset contains gene dependencies estimated for each gene and cell line using the CERES algorithm[22]. CRISPR KO gene dependency scores, which measures the sensitivity to the impact of CRISPR knockout-induced loss of a gene on cell viability, are normalized such that a value of 0 represents the median dependency score of negative control genes and −1 represents the median dependency score of sgRNAs that target pan-essential genes. CRISPR KO gene dependency scores were compared for cell lines with wild-type *PIK3CA* to cell lines harboring hotspot *PIK3CA* mutations and to cell lines with *PIK3CA* mutations observed in this AS cohort. *PIK3CA* hotspot mutations (as defined by depmap.org) were inferred using their frequency of occurrence in TCGA and COSMIC databases.

#### ***PIK3CA mutations in GENIE dataset***

The GENIE version 5 dataset was used to look for *PIK3CA* mutations and their occurrences in all cancer types[23] (http://genie.cbioportal.org/).
