## Supplemental Tables 1-7 for "The Angiosarcoma Project: enabling genomic and clinical discoveries in a rare cancer through patient-partnered research"

### Supplemental Table 1

#### Supplemental Table 1. Angiosarcoma Project Patient-Reported Data Summary (N=227)

| Question / Response | YES | NO | NOT REPORTED | DON'T KNOW | N/A |
|---|---|---|---|---|---|
| Are you currently being treated for your angiosarcoma? | 99 (43.6%) | 118 (52.0%) | 5 (2.2%) | 5 (2.2%) | -- |
| Have you had surgery to remove angiosarcoma? | 177 (78.0%) | 49 (21.6%) | 1 (0.4%) | 0 (0.0%) | -- |
| If so, did the surgery remove all known cancer tissue (also known as "clean margins")? | 142 (62.6%) | 30 (13.2%) | 1 (0.4%) | 5 (2.2%) | 49 (21.6%) |
| Have you had radiation as a treatment for angiosarcoma? If you had radiation for other cancers, we will ask you about that later. | 93 (41.0%) | 130 (57.3%) | 3 (1.3%) | 0 (0.0%) | 1 (0.4%) |
| Were you ever diagnosed with any other kind of cancer(s)? | 95 (41.9%) | 130 (57.3%) | 1 (0.4%) | 1 (0.4%) | -- |
| Have you had radiation as a treatment for another cancer(s)? | 60 (26.4%) | 164 (72.2%) | 1 (0.4%) | 1 (0.4%) | 1 (0.4%) |
| Do you consider yourself Hispanic, Latino/a or Spanish? | 9 (4.0%) | 214 (94.3%) | 4 (1.8%) | 0 (0.0%) | -- |

### Supplemental Table 2

**Supplemental Table 2. Patient reported sites of primary angiosarcoma. Number and percentage is indicated in the right column**

| Patient Reported Site of Primary Angiosarcoma | Number of Patients (%) |
|---|---|
| Breast (encompassing both cutaneous and parenchymal tumors of the breast) | 95 (42%) |
| Head, Face, Neck, Scalp | 56 (25%) |
| Bone/Limb | 21(9%) |
| Heart | 12 (5%) |
| Abdominal area | 6 (3%) |
| Liver | 4 (2%) |
| Spleen | 3 (1%) |
| Lung | 2 (1%) |
| Brain | 1 (0.4%) |
| Multiple locations | 16 (7%) |
| Other locations | 8 (3.5%) |
| No answer to the question | 3 (1%) |

### Supplemental Table 3

**Supplemental Table 3. Data Elements Abstracted from Medical Records of Sequenced Angiosarcoma Project Participants (N=36)**

| Patient-Level Data Elements | Sample-Level Data Elements |
|---|---|
| Biological Sex | Sample Collection Date |
| Age at Diagnosis | Procedure Location |
| Diagnositc Biopsy Location | Procedure Type |
| Primary Tumor Site | Vasoformative |
| Medications Received | Epithelioid |
| Radiation Associated Angiosarcoma | Spindle Cell |
| Adjuvant Radiation | Nuclear Grade |
| Presence of Metastatic Disease | |
| Time to Metastatic Diagnosis | |
| Metastatic Sites at Metastatic Diagnosis | |
| Ever Metastatic Sites | |
| Local Recurrence | |
| Local Recurrence Sites | |
| Time to Local Recurrence(s) | |
| Other Cancer Type | |

Obtained medical records contained sufficient information to assess and abstract all of the above data elements for 35 patients. For one additional patient, records were only sufficient to abstract 13 data elements. When available in medical records, dates of diagnoses and treatments were abstracted. To protect patient confidentiality, all dates reported in public releases are indicated as elapsed time relative to the date of initial angiosarcoma diagnosis. For additional details on medical record abstraction and data element masking, please see Supplemental Methods.

### Supplemental Table 4

#### Supplemental Table 4. *PIK3CA* Mutations Observed in Angiosarcoma Patients

| PI3K Protein Change | Patients with this mutation | Tumor fraction | Co-occurrence with other *PIK3CA* mutations in a patient | OncoKB Annotation | Cancer Hotspot | 3D Hotspot (as defined by Gao *et al.* 2017) | COSMIC occurrences | GENIE.v5 occurrences | Cancer cell lines (using DepMap) with observed mutation | Cancer cell types with functional validation *in vitro* | *In vitro* functional validation references | Other references | **Cumulative evidence of mutation being activating** |
|---|---|---|---|---|---|---|---|---|---|---|---|---|---|
| M1043I | ASCProject_loC8UQUk | 0.09 | T957P | Oncogenic, level_3b | Yes | No | 107 | 76 | | Breast, Colorectal | PMID: 26627007, PMID: 17376864, PMID: 15930273 | | Strong |
| M1043V | ASCProject_GXTMTxU7 | 0.23 | | Oncogenic, level_3b | Yes | No | 107 | 36 (multiple breast, multiple carcinomas) | | Breast | PMID: 26627007, PMID: 17376864 | | Strong |
| R88Q | ASCProject_3NflfGHo | 0.23 | | Likely Oncogenic, level_3b | Yes | Yes | 89 | 202 | 4 (2 *PIK3CA* mutation - soft tissue, uterus; 2 with other non-conserving *PIK3CA* mutations - colorectal, uterus) | Bone | PMID: 18829572 | PMID: 22949682 (Structural analysis) | Strong |
| N1044K | ASCProject_EAuOu4uD | 0.09 | | Oncogenic, level_3b | Yes | No | 32 | 35 (multiple breast, multiple carcinomas) | | Breast | PMID: 26627007 | | Strong |
| P124L | ASCProject_blh1uyhK | 0.33, 0.07 | | Unknown, level NA | No | No | 2 | 1 (Breast), 1(Angiosarcoma - P124Q), 1(Oligodendroglia - P124E) | 1 (urinary track) | Urothelial carcinoma | PMID: 22430209 PMID: 27465249 | | Medium |
| G914R | ASCProject_EdCVC0um | 0.06 | | Likely Oncogenic, level_3b | No | No | 1 | 8 (1 breast, 6 carcinomas) | 1(colorectal) | | PMID: 22729224, PMID: 27191687, PMID: 28502725 | | Medium |
| N107T | ASCProject_RvtOtjtj | 0.20, 0.14 | | Predicted Oncogenic, level_3b | No | No | 1 | 1 (Breast) | | | | PMID: 22949682 (Structural study on mutation in G106V) | Medium |
| T957P | ASCProject_loC8UQUk | 0.10, 0.20 | M1043I | Unknown, level NA | No | No | NA | 1 (Breast) | | | | | Unknown |

### Supplemental Table 5

#### Supplemental Table 5. Information on Angiosarcoma Patients Receiving Immunotherapy (N=6)

| Project ID | Primary Site | Tumor Mutation Burden (mutations per megabase) | Immunotherapy | IO Duration (days) | IO Response | Discontinuation | Received Tx After IO | Notes about IO Response |
|---|---|---|---|---|---|---|---|---|
| ASCProject_ bRSGSlCG | LUNG | 0.4 | PEMBROLIZUMAB | 42 | Varied response among lesions | Side Effects | YES | -- |
| ASCProject_ QYsAsrTO | PRIMARY BREAST | 4.6, 3.3 (two tumor samples) | NIVOLUMAB | 177 | Progression | Progression | YES | -- |
| | | | PEMBROLIZUMAB | 86 | Progression | Side Effects | UNKNOWN | Pembrolizumab was stopped due to adverse effects |
| ASCProject_ XjHAU9uR | CARDIAC | 1.4 | PEMBROLIZUMAB | 169 | Progression | Progression | YES | -- |
| ASCProject_ NdUxUwCM | HNFS | 62.3 | PEMBROLIZUMAB | 1 | Received one dose, progression | Progression, Side Effects | YES | Only 1 dose due to RA, clinical disease progression |
| ASCProject_ KxFGsofW | HNFS | 78.5 | PEMBROLIZUMAB | 219 | Complete | Side Effects | NO | Lasting response (NED) |
| ASCProject_ dyhLT8sG | HNFS | 138.9 | PEMBROLIZUMAB | 249; 30 | Stable disease | Side Effects | NO | Off therapy due to autoimmune hepatitis |

IO: Immunotherapy; Tx: Therapy (non-IO); HNFS: Head, Neck, Face and Scalp; RA: Rheumatoid Arthritis; NED: No Evidence of Disease

### Supplemental Table 6

Supplemental Table 6. Detailed timeline of all treatments received by two head, neck, face, scalp angiosarcoma patients who each had a complete response to Pembrolizumab. Number of days indicated below are calculated relative to the initial diagnosis of angiosarcoma (day of diagnosis being day 0).

| Patient ID | Start (Days) | Stop (Days) | Drug | Treatment Category |
|---|---|---|---|---|
| Patient 1 | | | | |
| ASCProject_KxFGsofW | 15 | 139 | PACLITAXEL | CHEMOTHERAPY |
| ASCProject_KxFGsofW | 15 | 148 | BEVACIZUMAB | VEGF INHIBITOR |
| ASCProject_KxFGsofW | 149 | 169 | NO TREATMENT | NA |
| ASCProject_KxFGsofW | 170 | 230 | GEMCITABINE | CHEMOTHERAPY |
| ASCProject_KxFGsofW | 170 | 230 | DOCETAXEL | CHEMOTHERAPY |
| ASCProject_KxFGsofW | 231 | 259 | NO TREATMENT | NA |
| ASCProject_KxFGsofW | 260 | 408 | DOXORUBICIN | CHEMOTHERAPY |
| ASCProject_KxFGsofW | 266 | 428 | DEXRAZOXANE | CHEMOPROTECTIVE AGENT |
| ASCProject_KxFGsofW | 300 | 408 | IFOSFAMIDE | CHEMOTHERAPY |
| ASCProject_KxFGsofW | 429 | 600 | NO TREATMENT | NA |
| ASCProject_KxFGsofW | 601 | 642 | CLINICAL TRIAL | CLINICAL TRIAL |
| ASCProject_KxFGsofW | 643 | 649 | NO TREATMENT | NA |
| ASCProject_KxFGsofW | 650 | 713 | CYCLOPHOSPHAMIDE | CHEMOTHERAPY |
| ASCProject_KxFGsofW | 714 | 747 | NO TREATMENT | NA |
| ASCProject_KxFGsofW | 748 | 814 | CLINICAL TRIAL | CLINICAL TRIAL |
| ASCProject_KxFGsofW | 814 | 833 | CLINICAL TRIAL | CLINICAL TRIAL |
| ASCProject_KxFGsofW | 833 | 860 | CRIZOTINIB | TYROSINE KINASE INHIBITOR |
| ASCProject_KxFGsofW | 860 | 1079 | PEMBROLIZUMAB | ANTI-PD-1 MAB |
| ASCProject_KxFGsofW | 1080 | 1847 | NO TREATMENT | NA |
| Patient 2 | | | | |
| ASCProject_dyhLT8sG | 0 | 450 | NO TREATMENT | NA |
| ASCProject_dyhLT8sG | 451 | 667 | PACLITAXEL PROTEIN BOUND | CHEMOTHERAPY |
| ASCProject_dyhLT8sG | 668 | 786 | NO TREATMENT | NA |
| ASCProject_dyhLT8sG | 787 | 907 | CLINICAL TRIAL | CLINICAL TRIAL |
| ASCProject_dyhLT8sG | 908 | 1133 | NO TREATMENT | NA |
| ASCProject_dyhLT8sG | 1134 | 1194 | PACLITAXEL PROTEIN BOUND | CHEMOTHERAPY |
| ASCProject_dyhLT8sG | 1195 | 1219 | NO TREATMENT | NA |
| ASCProject_dyhLT8sG | 1220 | 1469 | PEMBROLIZUMAB | ANTI-PD-1 MAB |
| ASCProject_dyhLT8sG | 1470 | 1681 | NO TREATMENT | NA |
| ASCProject_dyhLT8sG | 1682 | 1712 | PEMBROLIZUMAB | ANTI-PD-1 MAB |
| ASCProject_dyhLT8sG | 1713 | 2549 | NO TREATMENT | NA |

NA: Not Applicable, ANTI-PD-1 MAB: Anti-programmed-cell death-1 monoclonal antibody

Supplemental Table 7:

Angiosarcoma Project forms

Form 1. Blank copy of the Angiosarcoma Project primary screening questionnaire
Form 2. Blank copy of the Angiosarcoma Project patient intake survey
Form 3. Blank copy of the Angiosarcoma Project consent form
Form 4. Blank copy of the Angiosarcoma Project medical release form
Form 5. Blank copy of the medical record request form

#### 1. Blank copy of the Angiosarcoma Project primary screening questionnaire

Angiosarcoma Project

Learn More

### Join the movement: tell us about yourself

Complete the form below to tell us about yourself and your cancer. Our goal is to perform many different studies within the angiosarcoma community. Allowing us to know a little bit about your experience will help us conduct our current projects and also to design future studies. We will be starting with some focused studies in angiosarcoma, and expanding over time based on what we learn from you. We are asking all patients with angiosarcoma to say "Count Me In" and fill out the form so that we can use the information you provide to plan our next studies.

If you've previously submitted your name and email and have already started answering the questionnaire, click here so we can resend you a link that will let you pick up where you left off.

#### Contact Info

First Name *

Last Name *

Email Address *

Email Confirmation (Reenter Email) *

- ○ I have been diagnosed with angiosarcoma. I'm willing to answer additional questions about myself and my experience with angiosarcoma.
- ○ I haven't been diagnosed with angiosarcoma, but I want to stay informed about the Angiosarcoma Project by joining the email list.
- ○ I have a loved one that has passed away from angiosarcoma.

** Required field*

SUBMIT

Angiosarcoma Project

Home Data Release More Details/FAQ About Us Join Mailing List Information for Physicians ▲ Back to top

Count Me In

Contact Us:

857-500-6264

Count Me In
415 Main St, Cambridge, MA
02142, United States

**2. Blank copy of the Angiosarcoma Project patient intake survey**



Angiosarcoma Project

Learn More

### Join the movement: tell us about yourself

Thank you for providing your contact information. Please help us understand more about your angiosarcoma by answering the questions below.

As you fill out the questions below, your answers will be automatically saved. If you've previously entered information here and want to pick up where you left off, please use the link we sent you via email to return to this page.

If you would like your information deleted from our database, please let us know by emailing (mailto:) and we will remove your name and email address and the answers to any questions you may have answered.

#### About you

Please fill out as much as you can. All questions are optional. You can return at any time with the link sent to you by email.

1. When were you first diagnosed with angiosarcoma?

Choose month...
Angiosarcoma Project
Choose year...
Learn More

2. When you were <u>first</u> diagnosed with angiosarcoma, where in your body was it found (select all that apply)?

- ☐ Head/Face/Neck (not scalp)
- ☐ Scalp
- ☐ Breast
- ☐ Heart
- ☐ Liver
- ☐ Spleen
- ☐ Lung
- ☐ Brain
- ☐ Lymph Nodes
- ☐ Bone/Limb
- ☐ Abdominal Area
- ☐ Other

Please provide details

- ☐ I don't know

3. Please select all of the places in your body that you have <u>ever</u> had angiosarcoma (select all that apply).

- ☐ Head/Face/Neck (not scalp)
- ☐ Scalp
- ☐ Breast
- ☐ Heart

- [ ] Liver

Angiosarcoma Project

Learn More

- [ ] Spleen
- [ ] Lung
- [ ] Brain
- [ ] Lymph Nodes
- [ ] Bone/Limb
- [ ] Abdominal Area
- [ ] Other

Please provide details

- [ ] I don't know

4. Please select all of the places in your body where you currently have angiosarcoma (select all that apply). If you don't have evidence of disease, please select "No Evidence of Disease (NED)".

- [ ] Head/Face/Neck (not scalp)
- [ ] Scalp
- [ ] Breast
- [ ] Heart
- [ ] Liver
- [ ] Spleen
- [ ] Lung
- [ ] Brain
- [ ] Lymph Nodes
- [ ] Bone/Limb
- [ ] Abdominal Area

No Evidence of Disease (NED)

Angiosarcoma Project

Other

Learn More

Please provide details

I don't know

To help us understand the full scope of how your angiosarcoma was treated, the following questions will ask you separately about surgery, radiation, and any medications, drugs, or chemotherapies you may have received for angiosarcoma.

5. Have you had surgery to remove angiosarcoma?

- Yes
- No
- I don't know

If so, did the surgery remove all known cancer tissue (also known as "clean margins")?

- Yes
- No
- I don't know

6. Have you had radiation as a treatment for angiosarcoma? If you radiation for other cancers, we will ask you about that later.

- Yes
- No

☐ I don't know



Was your radiation before or after surgery?

- ☐ Before
- ☐ After
- ☐ Both
- ☐ I don't know

7. Please list the medications, drugs, and chemotherapies you have been prescribed specifically for the treatment of angiosarcoma. It’s okay if there are treatments you don’t remember. ⓘ

---

8. Are you currently being treated for your angiosarcoma?

- ☐ Yes
- ☐ No
- ☐ I don't know

Please list the therapies you are currently receiving for angiosarcoma (this can include upcoming surgeries, radiation, or medications, drugs, or chemotherapies).

---

9. Were you ever diagnosed with any other kind of cancer(s)?

- ☐ Yes
- ☐ No

☐ I don't know

Angiosarcoma Project

Learn More

Please list which cancer(s) and approximate year(s) of diagnosis.

Disease name ______ Year ______

+ADD ANOTHER CANCER

10. Have you had radiation as a treatment for another cancer(s)?

☐ Yes

☐ No

☐ I don't know

In what part of your body did you receive radiation for your other cancer(s)?

______

11. How did you hear about The Angiosarcoma Project?

______

12. Optional: Tell us anything else you want about yourself and your experience with angiosarcoma. We are asking this so you have an opportunity to tell us things that you feel are important for our understanding of this disease.

______

13. Do you consider yourself Hispanic, Latino/a or Spanish?

- [ ] Yes
- [ ] No
- [ ] I don't know

14. What is your race (select all that apply)?

- [ ] American Indian or Native American
- [ ] Japanese
- [ ] Chinese
- [ ] Other East Asian
- [ ] South East Asian or Indian
- [ ] Black or African American
- [ ] Native Hawaiian or other Pacific Islander
- [ ] White
- [ ] I prefer not to answer
- [ ] Other

15. In what year were you born?

Choose year...

16. What country do you live in?

Choose country...

17. What is your ZIP or postal code?

Zip Code

I [illegible]derstand that the information I entered here will be stored in a secure database a[illegible] may be used to match me to one or more research studies conducted by the A[illegible]osarcoma Project. If the information that I entered matches a study being conducted by the Angiosarcoma Project, either now or in the future, I agree to be contacted about possibly participating. I understand that if I would like my information deleted from the database, now or in the future, I can (mailto:) and my information will be removed from the database.

SUBMIT



Contact Us:
 (mailto:)

Count Me In
415 Main St, Cambridge, MA 02142, United States

##### 3. Blank copy of the Angiosarcoma Project consent form

BROAD INSTITUTE

415 Main Street
Cambridge, MA 02142
T 617-714-7000
www.broadinstitute.org

### Research Consent Form (Angiosarcoma Project)

Please read through the consent form text below and click Next when you are done to move on to the next section. If you have questions about the study or the consent form at any time, please contact us at the phone number or email address above.

**RESEARCH CONSENT FORM (Angiosarcoma Project) – KEY POINTS**

**"The Angiosarcoma Project"** is a patient-driven movement that empowers angiosarcoma patients to directly transform research and treatment of disease by sharing copies of their medical records and tissue and/or blood samples with researchers in order to accelerate the pace of discovery. Because we are enrolling participants across the country regardless of where they are being treated, this study will allow many more patients to contribute to research than has previously been possible.

#### 1. What is the purpose of this study?

We want to understand angiosarcoma better so that we can develop more effective therapies. By partnering directly with patients, we are able to study many more aspects of cancer than would otherwise be possible.

#### 2. What will I have to do if I agree to participate in this study?

Participation requires little effort. With your permission, we may ask that you send a saliva sample to us in a pre-stamped package that we will provide. We may also ask for your medical records. If so, we will take care of obtaining copies of your medical records from the hospitals or centers where you receive your medical care. If you elect to share tissue with us, we may also obtain small amounts of your stored tumor tissues from hospitals or centers where you receive your care. If you elect to share blood with us, we may ask you to have a sample of blood (1 tube or 2 teaspoons) drawn at your physician's office, local clinic, or nearby lab facility – we will provide detailed instructions on how to do this.

#### 3. Do I have to participate in this study?

No. Taking part in this study is voluntary. Even if you decide to participate, you can always change your mind and leave the study.

415 Main Street
Cambridge, MA 02142
T 617-714-7000
www.broadinstitute.org

#### 4. Will I benefit from participating?

While taking part in this study may not improve your own health, the information we collect will aid in our research efforts to provide better cancer treatment and prevention options to future patients. We will provide updates about key research discoveries made possible by your participation on our website.

#### 5. What are the risks of taking part in this research?

If you elect to share blood, there are small risks associated with obtaining a sample of blood. You may experience slight pain and swelling at the site of the blood draw. These complications are rare and should resolve within a few days. If they do not, you should contact your doctor. There may be a risk that your information (which includes your genetic information and information from your medical records) could be seen by unauthorized individuals. However, we have procedures and security measures in place designed to minimize this risk and protect the confidentiality of your information.

#### 6. Will it cost me anything to participate in this study?

No.

#### 7. Who will use my samples and see my information?

Your samples and health information will be available to researchers at the Broad Institute of MIT and Harvard, a not-for-profit biomedical research institute. After removing your name and other readily identifiable information, we will share results obtained from your participation with the greater research community as well as central data banks at the National Institutes of Health.

#### 8. Can I stop taking part in this research study?

Yes, you can withdraw from this research study at any time, although any of your information that has already been entered into our system cannot be withdrawn. Your information would be removed from future studies.

#### 9. What if I have questions?

415 Main Street
Cambridge, MA 02142
T 617-714-7000
www.broadinstitute.org

### FULL RESEARCH CONSENT FORM

#### The Angiosarcoma Project

##### A. Introduction

You are being invited to participate in a research study that will collect and analyze samples and health information of patients with angiosarcoma. This study will help doctors and researchers better understand why angiosarcoma occurs and develop ways to better treat and prevent it.

Cancers occur when the molecules that control normal cell growth (genes and proteins) are altered. Changes in the genes of tumor cells and normal tissues are called “alterations.” Several alterations that occur in certain types of cancers have already been identified and have led to the development of new drugs that specifically target those alterations. However, the vast majority of tumors from patients have not been studied, which means there is a tremendous amount of information still left to be discovered. Our goal is to discover more alterations, and to better understand those that have been previously described. We think this could lead to the development of additional therapies and cures.

Genes are composed of DNA “letters,” which contain the instructions that tell the cells in our bodies how to grow and work. We would like to use your DNA to look for alterations in cancer cell genes using a technology called “sequencing.”

Gene sequencing is a way of reading the DNA to identify alterations in genes that may contribute to the behavior of cells. Some changes in genes occur only in cancer cells. Others occur in normal cells as well, in the genes that may have been passed from parent to child. This research study will examine both kinds of genes.

You are being asked to participate in the study because you have angiosarcoma. Other than providing samples of saliva and, if you elect to, blood (1 tube or 2 teaspoons), participating in the study involves no additional tests or procedures.

This form explains why this research study is being done, what is involved in participating, the possible risks and benefits of the study, alternatives to participation, and your rights as a participant. The decision to participate is yours. We encourage you to ask questions about the study now or in the future.

##### B. Why is this research study being done?

We want to understand cancer better so that we can develop more effective therapies. By partnering directly with patients, we will be able to study many more aspects of cancer than has previously been possible. In addition, because we are enrolling participants across the country regardless of where they are being treated, this study will allow many more patients to directly contribute to research than might otherwise be feasible.

BROAD INSTITUTE

415 Main Street
Cambridge, MA 02142
T 617-714-7000
www.broadinstitute.org

#### C. What other options are there?

Taking part in this research study is voluntary – you may choose not to participate. Your decision not to participate will not affect your medical care in any way or result in any penalty or loss of benefits.

#### D. What is involved in the research study?

With your consent, we may obtain copies of your medical records and ask that you collect a sample of saliva at home–we will provide detailed instructions on how to do this. If you elect to share tissue samples with us, we may request a portion of your tumor tissues through already stored biopsies or surgical specimens in hospitals or centers where you received your medical care in the past. If you elect to share blood samples with us as well, we may ask you to have a sample of blood (1 tube or 2 teaspoons) drawn at your physician's office, local clinic, or nearby lab facility. We'll ask you to send any blood and/or saliva sample(s) to us in pre-stamped packages that we will provide.

We will analyze the genes in your cancer cells (obtained from the sample of your blood) and your normal cells (obtained from the sample of your blood or from your saliva sample). No additional procedures will be required. The results of this analysis will be used to try to develop better ways to treat and prevent cancers.

We will link the results of the gene tests on your cancer cells and normal cells with medical information that has been generated during the course of your treatment. We are asking your permission to obtain a copy of your medical record from places where you have received care for your cancer.

In some cases, a research doctor may contact you to find out if you would be interested in participating in a different or future research study based on information that may have been found in your samples.

To allow sharing of information with other researchers, the National Institutes of Health (NIH) and other organizations have developed central data (information) banks that analyze information and collect the results of certain types of genetic studies. These central banks will store your genetic and medical information and provide the information to qualified researchers to do more studies. We will also store your genetic and medical information at the Broad Institute of MIT and Harvard and share your information with other qualified researchers. Therefore, we are asking your permission to share your results with these special banks and other researchers, and have your information used for future research studies, including studies that have not yet been designed, studies involving diseases other than cancer, and/or studies that may be for commercial purposes (such as the development or approval of new drugs). Your information will be sent to central banks and other researchers only with a code number attached. Your name, social security number, and other information that could readily identify you will not be shared with central banks or other researchers. We will never sell your readily identifiable information to anyone under any circumstances.

#### E. How long will I be in this research study?

You may be asked to give samples of blood, tissue, and/or saliva after you consent to enrolling in this study. We will keep your blood, tissue, and saliva samples and medical records indefinitely until this

415 Main Street
Cambridge, MA 02142
T 617-714-7000
www.broadinstitute.org

study is finished, unless you inform us that you no longer wish to participate. You may do this at any time. More information about how to stop being in the study is below in paragraph I.

Once the study is finished, any left over blood and saliva samples and your medical records will be destroyed. Any tissue samples that we have will be returned to the pathology department at the hospital or other place where you received treatment.

#### F. What kind of information could be found in this study and will I be able to see it?

The gene tests in this study are being done to add to our knowledge of how genes and other factors affect cancer. This information will be kept confidential and while you will not receive information about your personal results obtained from studying your blood, saliva, or tissue samples, we will provide general results and major discoveries to all participants. We will do this by regularly updating the website that you used to enroll in this study. Furthermore, we will publish important discoveries found through these studies in the scientific literature so that the entire research community can work together to better understand cancer. Your individual data will not be published in a way in which you could be readily identified. Abstracts, which are plain language summaries of the published reports, will be available to you and the general public.

#### G. What are the risks or discomforts of the research study?

If you elect to share blood, there are small risks associated with obtaining the tube of blood. You may experience slight pain and swelling at the site of the blood draw. These complications are rare and should resolve within a few days. If they do not, you should contact your doctor.

There is a small risk that by participating in this study, the gene test results, including the identification of genetic changes in you or your cancer, could be seen by unauthorized individuals. We have tried to minimize this risk by carefully limiting access to the computers that would house your information to the staff of this research study.

There is a small but real risk that if your samples are used for this research study, they might not be available for clinical care in the future. However, we have attempted to minimize this risk in the following way: the pathologists in the department of pathology where your specimens are kept will not release your specimen unless they believe that the material remaining after the research test is performed is sufficient for any future clinical needs.

#### H. What are the benefits of the research study?

Taking part in this research study may not directly benefit you. By joining this study, you will help us and other researchers understand how to use gene tests to improve the care of patients with cancer in the future. We will provide study participants updates on our project website about key research discoveries made possible by your participation

#### I. Can I stop being in the research study and what are my rights?

You can stop being in the research study at any time. We will not be able to withdraw all the information that already has been used for research. If you tell us that you want to stop being in the study, we will return any remaining tumor samples from where we obtained them, and destroy any remaining blood, saliva samples, or DNA samples we have. We will not perform any additional tests on the samples. Additionally, we will not collect any additional medical records and we will destroy the medical records we already have. However, we will keep the results from the tests we did before you stopped being in the study. We will also keep the information we learned from reviewing your medical records before you stopped being in the study, We will not be able to take back the information that already has been used or shared with other researchers, central data banks, or that has been used to carry out related activities such as oversight, or that is needed to ensure quality of the study.

To withdraw your permission, you must do so in writing by contacting the researcher listed below in the section: "Whom do I contact if I have questions about the research study?" If you choose to not participate, or if you are not eligible to participate, or if you withdraw from this research study, this will not affect your present or future care and will not cause any penalty or loss of benefits to which you are otherwise entitled.

#### J. Will I be paid to take part in this research study?

There is no financial compensation for participation in this study.

#### K. What are the costs?

There are no costs to you to participate in this study.

#### L. What happens if I am injured or sick because I took part in this research study?

There is little risk that you will become injured or sick by taking part in this study. There are no plans for this project to pay you or give you other compensation for any injury. You do not give up your legal rights by signing this form. If you think you have been injured as a result of taking part in this research study, please tell the person in charge of this research study as soon as possible. The research doctor's contact information is listed in this consent form.

#### M. What about confidentiality?

We will take rigorous measures to protect the confidentiality and security of all your information, but we are unable to guarantee complete confidentiality. Information shared with the research team through email, or information accessible from a link in an email, is only protected by the security measures in place for your email account. Information from your medical records and genomics tests will be protected in a HIPAA compliant database.

When we receive any of your samples, your name, social security number, and other information that could be used to readily identify you will be removed and replaced by a code. If we send your samples to our collaborators for gene testing, the samples will be identified using only this code. The medical records that we receive will be reviewed by our research team to confirm that you are eligible for the study and to obtain information about your medical condition and treatment.

We will store all of your identifiable information related to the study (including your medical records) in locked file cabinets and in password-protected computer files at the Broad Institute and we will limit access to such files. We may share your identifiable information or coded information, as necessary, with regulatory or oversight authorities (such as the Office for Human Research Protections), ethics committees reviewing the conduct of the study, or as otherwise required by law.

When we send the results of the gene tests and your medical information to central data banks or other researchers, they will not contain your name, social security number, or other information that could be used to readily identify you.

The results of this research study or future research studies using the information from this study may be published in research papers or included in presentations that will become part of the scientific literature. You will not be identified in publications or presentations.

#### N. Whom do I contact if I have questions about the research study?

- Nikhil Wagle, MD
- Corrie Painter, PhD

For questions about your rights as a patient, please contact a representative of the Office for Human Research Studies at (617) 632-3029. This can include questions about your participation in the study, concerns about the study, a research related injury, or if you feel/felt under pressure to enroll in this research study or to continue to participate in this research study. Please keep a copy of this document in case you want to read it again.

#### O. Authorization to use your health information for research purposes

Because information about you and your health is personal and private, it generally cannot be used in this research study without your written authorization. Federal law requires that your health care providers and healthcare institutions (hospitals, clinics, doctor's offices) protect the privacy of information that identifies you and relates to your past, present, and future physical and mental health conditions.

If you sign this form, it will provide your health care providers and healthcare institutions the authorization to disclose your protected health information to the Broad Institute for use in this research study. The form is intended to inform you about how your health information will be used or disclosed in the study. Your information will only be used in accordance with this authorization form and the informed consent form and as required or allowed by law. Please read it carefully before signing it.

415 Main Street
Cambridge, MA 02142
T 617-714-7000
www.broadinstitute.org

1. **What personal information about me will be used or shared with others during this research?**
   - Health information created from study-related tests and/or questionnaires
   - Your medical records
   - Your saliva sample

   **If elected (at the end of this form):**
   - Your blood sample
   - Your tissue samples relevant to this research study and related records
2. **Why will protected information about me be used or shared with others?**

   The main reasons include the following:
   - To conduct and oversee the research described earlier in this form;
   - To ensure the research meets legal, institutional, and accreditation requirements;
   - To conduct public health activities (including reporting of adverse events or situations where you or others may be at risk of harm)
3. **Who will use or share protected health information about me?**

   The Broad Institute and its researchers and affiliated research staff will use and/or share your personal health information in connection with this research study.
4. **With whom outside of the Broad Institute may my personal health information be shared?**

   While all reasonable efforts will be made to protect the confidentiality of your protected health information, it may also be shared with the following entities:
   - Federal and state agencies (for example, the Department of Health and Human Services, the Food and Drug Administration, the National Institutes of Health, and/or the Office for Human Research Protections), or other domestic or foreign government bodies if required by law and/or necessary for oversight purposes. A qualified representative of the FDA and the National Cancer Institute may review your medical records.
   - Outside individuals or entities that have a need to access this information to perform functions relating to the conduct of this research such as data storage companies.

   Some who may receive your personal health information may not have to satisfy the privacy rules and requirements. They, in fact, may share your information with others without your permission.
5. **For how long will protected health information about me be used or shared with others?**

   There is no scheduled date at which your protected health information that is being used or shared for this research will be destroyed, because research is an ongoing process.
6. **Statement of privacy rights:**
   - You have the right to withdraw your permission for the doctors and researchers to use or share your protected health information. We will not be able to withdraw all the information that already

has been used or shared with others to carry out related activities such as oversight, or that is needed to ensure quality of the study. To withdraw your permission, you must do so in writing by contacting the researcher listed above in the section: "Whom do I contact if I have questions about the research study?"

- You have the right to request access to your personal health information that is used or shared during this research and that is related to your treatment or payment for your treatment. To request this information, please contact your doctor who will request this information from the study directors.

#### P. Participation Information

If you decide to sign this consent form, we may ask you for information about contacting your physicians and the hospitals that you were treated at for your cancer. We will not disclose details about the results of your participation in this study with any of the individuals that we contact, but rather ask them to provide us with your medical history and your tissue samples.

#### Q. Documentation of Consent

**This is what I agree to:**

**Please Check "Yes" or "No" for each point below:**

- You can work with me to arrange a sample of blood to be drawn at my physician's office, local clinic, or nearby lab facility. ☐ **Yes** ☐ **No**
- You can request my stored tissue samples from my physicians and the hospitals and other places where I received my care, perform (or collaborate with others to perform) gene tests on the samples, and store the samples until this research study is complete. ☐ **Yes** ☐ **No**

**In addition, I agree to all of the following:**

- You can request my medical records from my physicians and the hospitals and other places where I received and/or continue to receive my treatment and link results of the gene tests you perform on my saliva and, if I elect on this form, blood and tissue samples with my medical information from my medical records.
- You can analyze a saliva sample that I will send you, link the results to my medical information and other specimens, and store the specimen to use it for future research.
- You can perform (or collaborate with others to perform) gene tests on the blood and saliva samples that I will send you and store the samples until this research study is complete.
- You can use the results of the gene tests and my medical information for future research studies, including studies that have not yet been designed, studies for diseases other than cancer, and/or studies that may be for commercial purposes.
- You can share the results of the gene tests and my medical information with central data banks (e.g., the NIH) and with other qualified researchers in a manner that does not include my name,

415 Main Street
Cambridge, MA 02142
T 617-714-7000
www.broadinstitute.org

social security number, or any other information that could be used to readily identify me, to be used by other qualified researchers to perform future research studies, including studies that have not yet been designed, studies for diseases other than cancer, and studies that may be for commercial purposes.

**My full name below indicates:**

- ❑ I have had enough time to read the consent and think about agreeing to participate in this study;
- ❑ I have had all of my questions answered to my satisfaction;
- ❑ I am willing to participate in this research study;
- ❑ I have been told that my participation is voluntary and if I decide not to participate it will have no impact on my medical care;
- ❑ I have been told that if I decide to participate now, I can decide to stop being in the study at any time.
- ❑ I acknowledge that a copy of the signed consent form will be sent to my email address

Your Full Name:

________________________________________

Date of Birth (mm/dd/yyyy)

__________________

Date

__________________________

03/29/194 __________________________

#### 4. Blank copy of the Angiosarcoma Project medical release form

BROAD INSTITUTE

415 Main Street
Cambridge, MA 02142
T 617-714-7000
www.broadinstitute.org

**Thank you very much for your consent to participate in this research study. To complete the process, we will need to collect some additional information from you below:**

**To proceed with this study, we need to collect information about:**

1. **Your contact information, including your current mailing address, so that we can send you a saliva kit**
2. **The name and contact information for the physician(s) who has/have cared for you throughout your experiences with angiosarcoma, so we can obtain copies of your medical records**
3. **The names of the hospitals / institutions where you've had biopsies and surgeries, so we can obtain some of your stored tumor samples**

**Printed below is the information you have provided to us:**

**YOUR CONTACT INFORMATION:**

**First Name:** ________________________________________

**Last Name:** ________________________________________

**Current Mailing Address:** ________________________________________

**City:**_____________________**State:**_____________________**Zip:** _______________**Country:** _____

**Phone:** ________________________________________

BROAD INSTITUTE

415 Main Street
Cambridge, MA 02142
T 617-714-7000
www.broadinstitute.org

#### YOUR PHYSICIANS' NAMES:

Physician Name: ___________________________________________

Institution (if any): ___________________________________________

City: _______________________State:_______________________

415 Main Street
Cambridge, MA 02142
T 617-714-7000
www.broadinstitute.org

**YOUR INITIAL BIOPSY HOSPITAL / INSTITUTION NAME:**

**Institution:** ____________________________________________

**City:**______________________________**State:**____________________

415 Main Street
Cambridge, MA 02142
T 617-714-7000
www.broadinstitute.org

**By completing this information, you are agreeing to allow us to contact these physician(s) and hospital(s) / institution(s) to obtain your records.**

- **I have already read and signed the informed consent document for this study, which describes the use of my personal health information (Section O), and hereby grant permission to Nikhil Wagle, MD, Dana-Farber Cancer Institute, 450 Brookline Ave, Boston, MA, 02215, or a member of the study team to examine copies of my medical records pertaining to my angiosarcoma diagnosis and treatment, and, if I elected on the informed consent document, to obtain tumor tissue for research studies. I acknowledge that a copy of this completed form will be sent to my email address.**

______________________ Full Name

______________________ Date

______________________ Date of Birth

**5. Blank copy of the medical record request form**

**Nikhil Wagle, MD**
Broad Institute
ASCProject, Cancer Program
415 Main Street, Room 4041
Cambridge, MA 02142
P: 617-714-8510 F: 617-395-2631
E:

### Fax

**To:**
**Fax:**

From: Nikhil Wagle, MD
Phone Number: 617-714-8510
Fax Number: 617-395-2631

Number of Pages Including Cover Page: # 19
Date:

**NOTES:** The following materials are enclosed for a patient enrolled on DFCI Protocol 15-057, and we are requesting the patient's medical records as part of the study procedure. The following items are enclosed as part of our request.

1. Medical Record Request Letter (i.e., outline of requested materials)
2. Electronically-signed patient consent form
3. Electronically-signed Medical Record Release Form
4. Latest IRB approval memo

**IMPORTANT**: Please note that consent & release forms **DO NOT EXPIRE**. Participants sign consent and complete the medical record release form only once when they are enrolled. Participants are never re-consented for this minimal risk study and consent forms remain valid with annual study renewal per IRB approval (included). Please call Rachel Stoddard, the study's Clinical Research Coordinator, at 617-714-8510 if you have questions.

**Confidential**

The documents accompanying this fax transmission may contain confidential patient information belonging to the sender that is legally privileged. This authorized recipient of this information is prohibited from disclosing this information to any other party. If you have received this transmission in error, please notify the sender immediately. Thank you.

Dear

I am writing to request the medical records for the following patient:

- Patient Name:
- DOB:
- Diagnosis Date:

We are reviewing this patient's medical records as part of their participation in *The Angiosarcoma Project* at Dana-Farber Cancer Institute and the Broad Institute of MIT and Harvard. We will collect medical record data on an as needed basis from participants' physicians.

We are requesting the following documents from the patients' medical record:

- All clinic notes from treating providers, including medical oncologists, residents, fellows, radiation oncologists, surgeons, nurse practitioners, etc. **from the date of diagnosis through present day ______________.**
- Angiosarcoma treatment data (including radiation, chemotherapy and hormonal therapy).
- Pathology reports
- Operative reports
- Medications, pharmacy records & infusion center records
- Referrals
- MD to MD exchange
- Genetic testing reports

Please find the patients' electronically-signed and dated Research Consent Form, as well as the Patient Release of Medical Records form also electronically-signed by the patient, accompanying this letter.

**Nikhil Wagle, MD**
**Broad Institute**
**ASCProject, Cancer Program**
**415 Main Street, Room 4041**
**Cambridge, MA 02142**

If you have any questions, please feel free to contact Clinical Research Coordinator Rachel Stoddard at 617-714-8510.

Thank you very much for your help with our study!

**Confidential**

The documents accompanying this fax transmission may contain confidential patient information belonging to the sender that is legally privileged. This authorized recipient of this information is prohibited from disclosing this information to any other party. If you have received this transmission in error, please notify the sender immediately. Thank you.
