## Supplemental Figures for "The Angiosarcoma Project: enabling genomic and clinical discoveries in a rare cancer through patient-partnered research"

### Supplemental Figure 1

#### 340 Institutions Represented Among 225 US & Canadian Patients

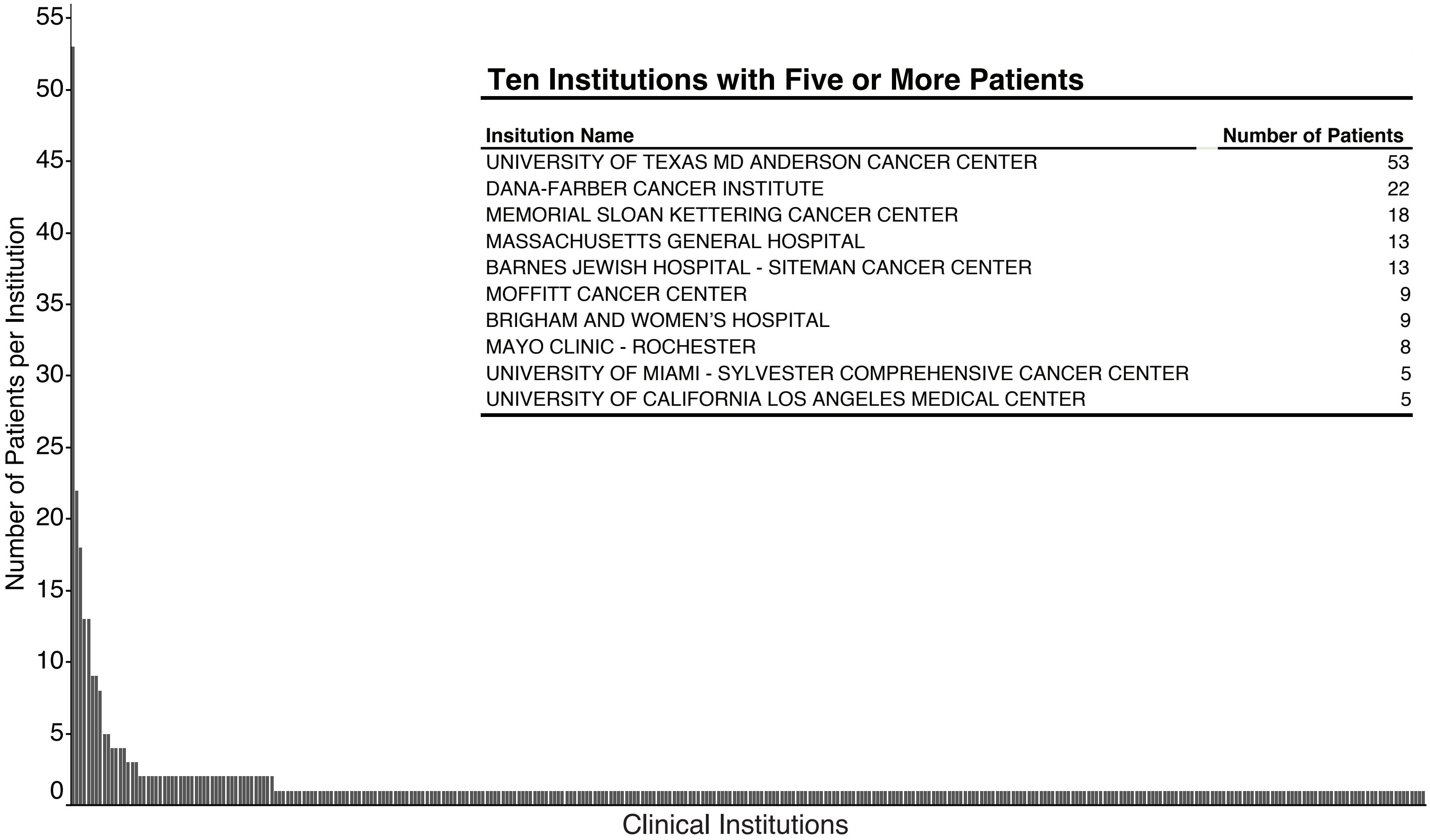


**Ten Institutions with Five or More Patients**

| Insitution Name | Number of Patients |
|---|---|
| UNIVERSITY OF TEXAS MD ANDERSON CANCER CENTER | 53 |
| DANA-FARBER CANCER INSTITUTE | 22 |
| MEMORIAL SLOAN KETTERING CANCER CENTER | 18 |
| MASSACHUSETTS GENERAL HOSPITAL | 13 |
| BARNES JEWISH HOSPITAL - SITEMAN CANCER CENTER | 13 |
| MOFFITT CANCER CENTER | 9 |
| BRIGHAM AND WOMEN'S HOSPITAL | 9 |
| MAYO CLINIC - ROCHESTER | 8 |
| UNIVERSITY OF MIAMI - SYLVESTER COMPREHENSIVE CANCER CENTER | 5 |
| UNIVERSITY OF CALIFORNIA LOS ANGELES MEDICAL CENTER | 5 |

Supplemental Figure 1. Clinical instituions at which Angiosarcoma Project patients received care for AS. This histogram depicts the 340 different clinical institutions that ASCproject patients reported they received some aspect of their clinical care for AS (x-axis) and the number of patients that reported care at any given institution (y-axis). This information was provided on the medical release forms completed by consented Angiosarcoma Project patients (n=225 patients). Of the 227 consented ASCProject participants, 2 people did not fill out the medical release form, and are not included in this figure. The majority of the 340 total institutions were listed by only one patient. Ten institutions were listed by five or more AS patients as sites of care (box, upper right).
AS, angiosarcoma; ASCproject, The Angiosarcoma Project.

### Supplemental Figure 2

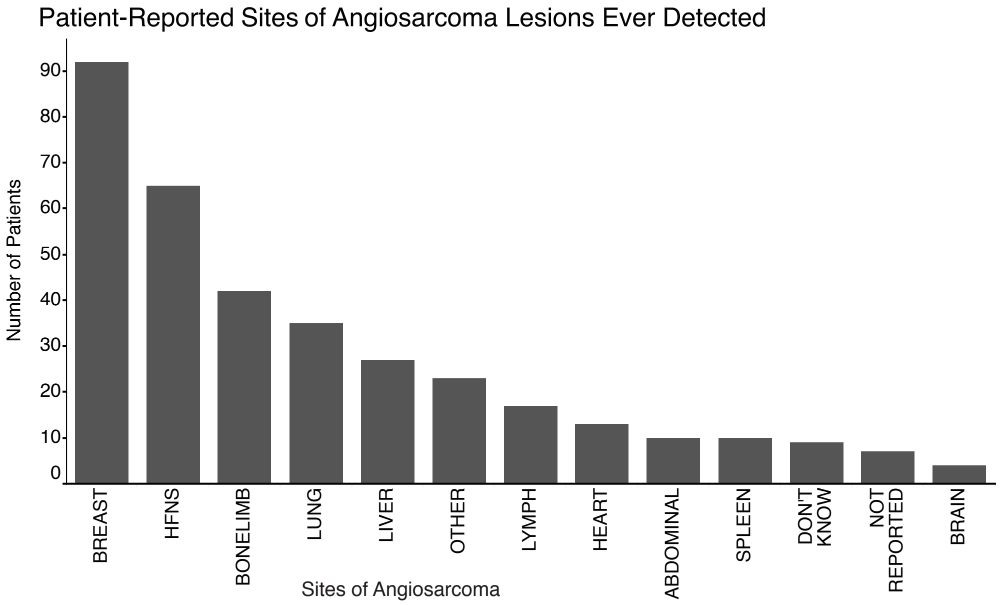


Supplemental Figure 2. Patient-reported sites of angiosarcoma lesions ever detected. A histogram showing the patient responses to the intake survey question ‘Please select all of the places in your body that you have ever had angiosarcoma (Please select all that apply).’ Patients first complete the patient intake survey during the Angiosarcoma Project registration process. Patient intake surveys completed by the 227 patients who consented for the Angiosarcoma Project as of September 30, 2018 were analyzed. Each response of a location is counted separately, such that multiple locations are shown for patients that selected more than one answer to indicate that they have had more than one location for their angiosarcoma lesions. 9 patients selected the provided response option of ‘I don’t know’ (‘Don’t Know’), and 7 patients did not respond to this question (’Not Reported’). The HNFS category depicted above includes patient responses of ‘head, face, neck’ and responses of ‘scalp.’ HNFS, head, neck, face, scalp.

### Supplemental Figure 3

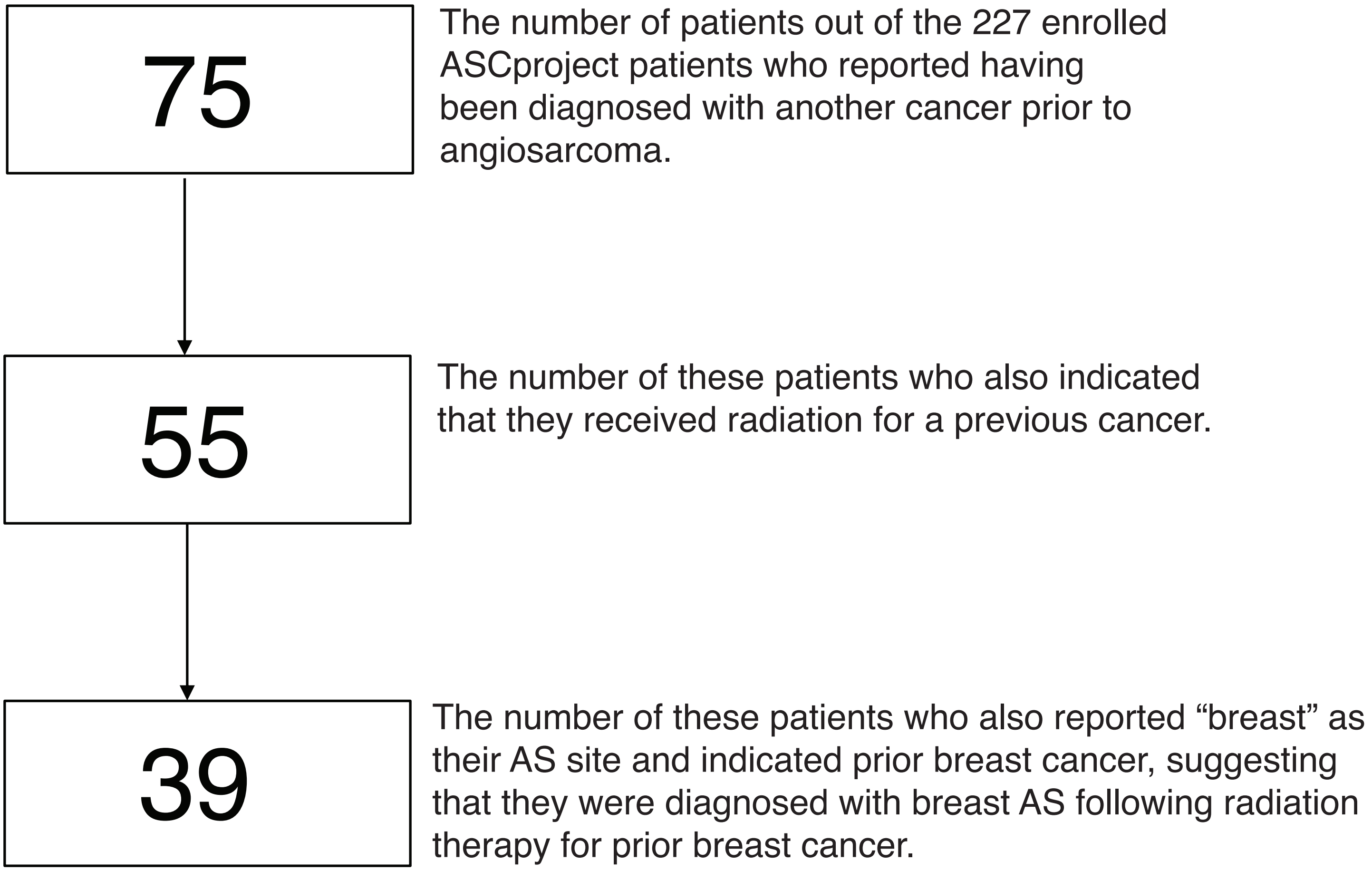


Supplemental Figure 3. Patient-reported information regarding other cancer diagnoses prior to angiosarcoma. Patient intake surveys completed by the 227 patients who consented for the Angiosarcoma Project as of September 30, 2018 were analyzed. Seventy five patients' responses indicated they had been diagnosed with another cancer prior to angiosarcoma. These patients were identified based on their responses of "Yes" to the survey question "Were you ever diagnosed with any other kind of cancer(s)?". Patients also provided specific diagnoses years for AS and other cancer(s), which were used to determine that the AS diagnosis occurred in the same year or after another cancer diagnosis. Of those 75 patients, 55 also responded "Yes" to the survey question "Have you had radiation as a treatment for another cancer(s)?". Of those 55 patients, 39 also reported “breast” as their only primary AS site and indicated breast cancer as a previous cancer. The majority of these 39 patients are expected to be cases of cutaneous AS of the breast.
AS, Angiosarcoma; ASCproject, The Angiosarcoma Project

### Supplemental Figure 4

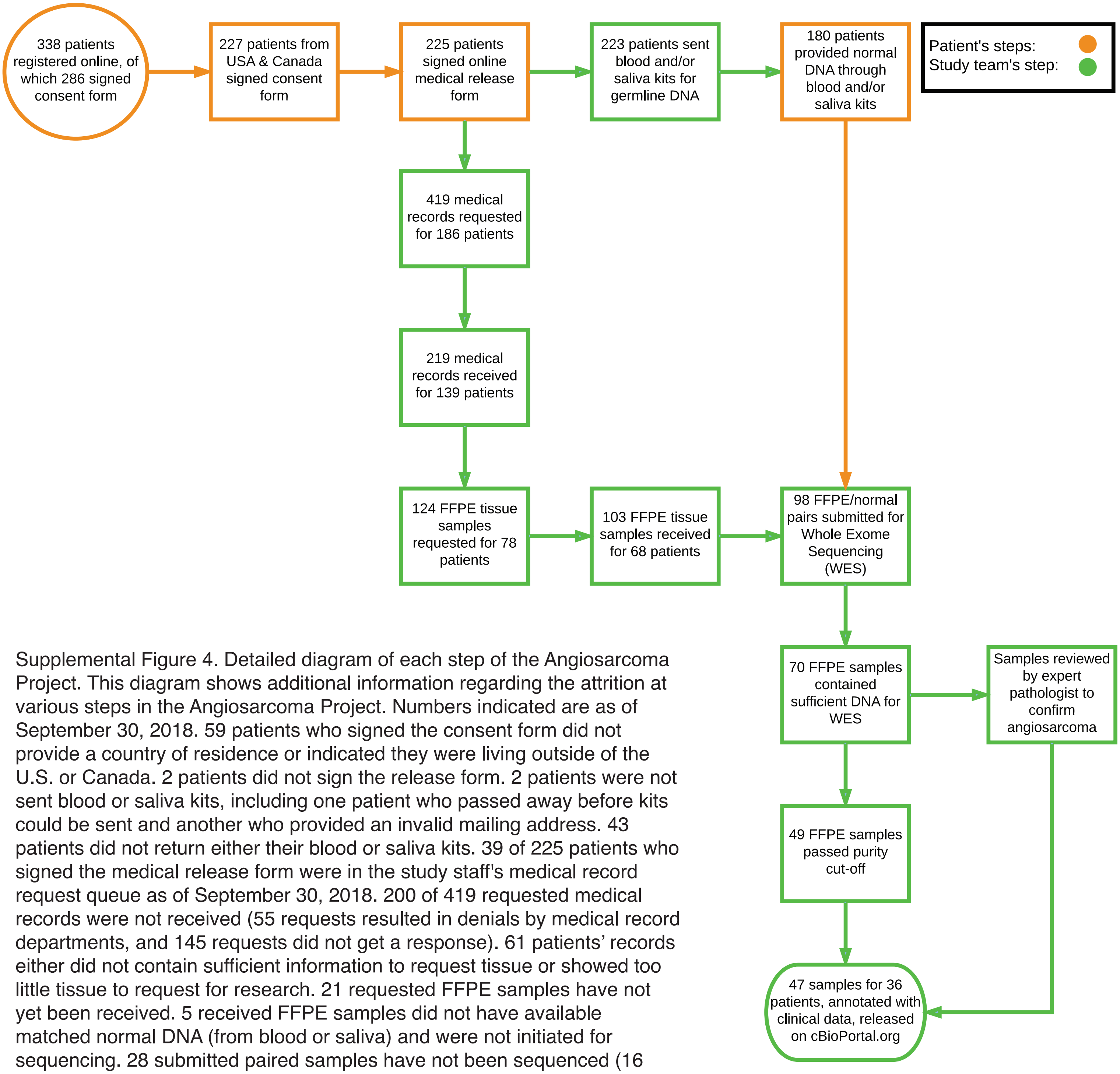


Supplemental Figure 4. Detailed diagram of each step of the Angiosarcoma Project. This diagram shows additional information regarding the attrition at various steps in the Angiosarcoma Project. Numbers indicated are as of September 30, 2018. 59 patients who signed the consent form did not provide a country of residence or indicated they were living outside of the U.S. or Canada. 2 patients did not sign the release form. 2 patients were not sent blood or saliva kits, including one patient who passed away before kits could be sent and another who provided an invalid mailing address. 43 patients did not return either their blood or saliva kits. 39 of 225 patients who signed the medical release form were in the study staff's medical record request queue as of September 30, 2018. 200 of 419 requested medical records were not received (55 requests resulted in denials by medical record departments, and 145 requests did not get a response). 61 patients' records either did not contain sufficient information to request tissue or showed too little tissue to request for research. 21 requested FFPE samples have not yet been received. 5 received FFPE samples did not have available matched normal DNA (from blood or saliva) and were not initiated for sequencing. 28 submitted paired samples have not been sequenced (16 samples had insufficient material for sequencing and 12 sets of samples are still in the sequencing pipeline). 21 sequenced samples had less than 10% tumor purity. 2 samples were excluded from dataset because centralized pathology review determined the samples were not angiosarcoma. The remaining 47 FFPE samples from 36 patients that underwent whole exome sequencing comprised the Angiosarcoma Project September 2018 dataset that was released on cBioPortal.org along with associated patient-reported and clinical data.

### Supplemental Figure 5

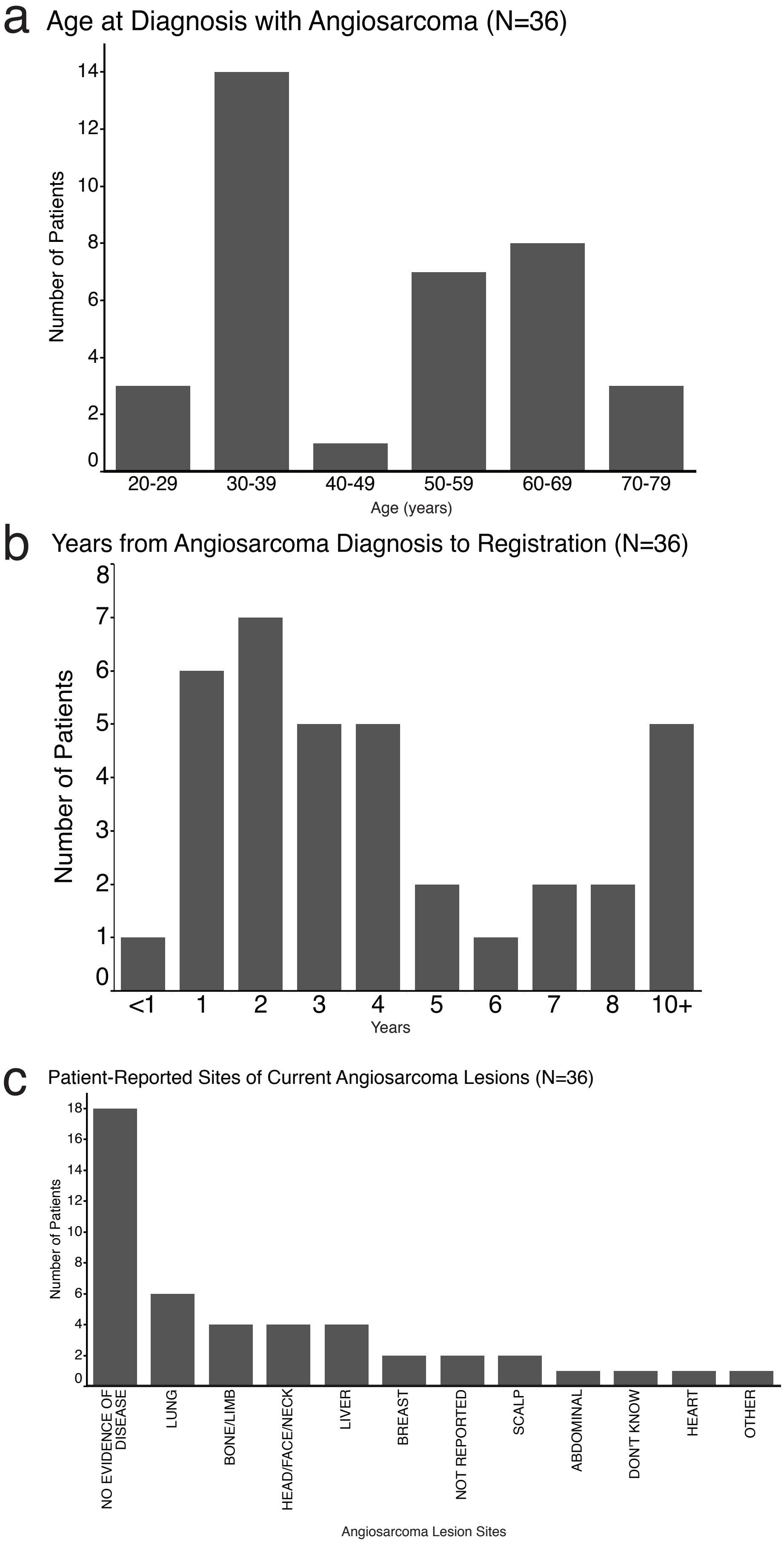


Supplemental Figure 5. Patient-reported data of 36 AS patients whose samples were sequenced in the Angiosarcoma Project. The intake survey completed during the ASCproject registration process by these 36 patients were analyzed. (a) A histogram showing the age in years of these 36 patients at initial diagnosis with AS (mean: 47.8 years). These values were calculated from patient provided date of birth and date of initial AS diagnosis. (b) A histogram showing the years elapsed between these 36 patients' initial diagnosis with AS and patients' registration in the ASCproject (mean: 4.6 years). These values were calculated from the date of project registration and the patient-provided date of initial AS diagnosis. (c) A histogram showing the patient-reported location of angiosarcoma at the time of last intake survey completion. An option was provided for patients to report no evidence of disease. Patients with more than one location of AS were able to provide more than one site. 2 patients did not respond to this question ('Not Reported') and 1 patient responded 'Don't Know.'
AS, Angiosarcoma; ASCproject, The Angiosarcoma Project.

### Supplemental Figure 6

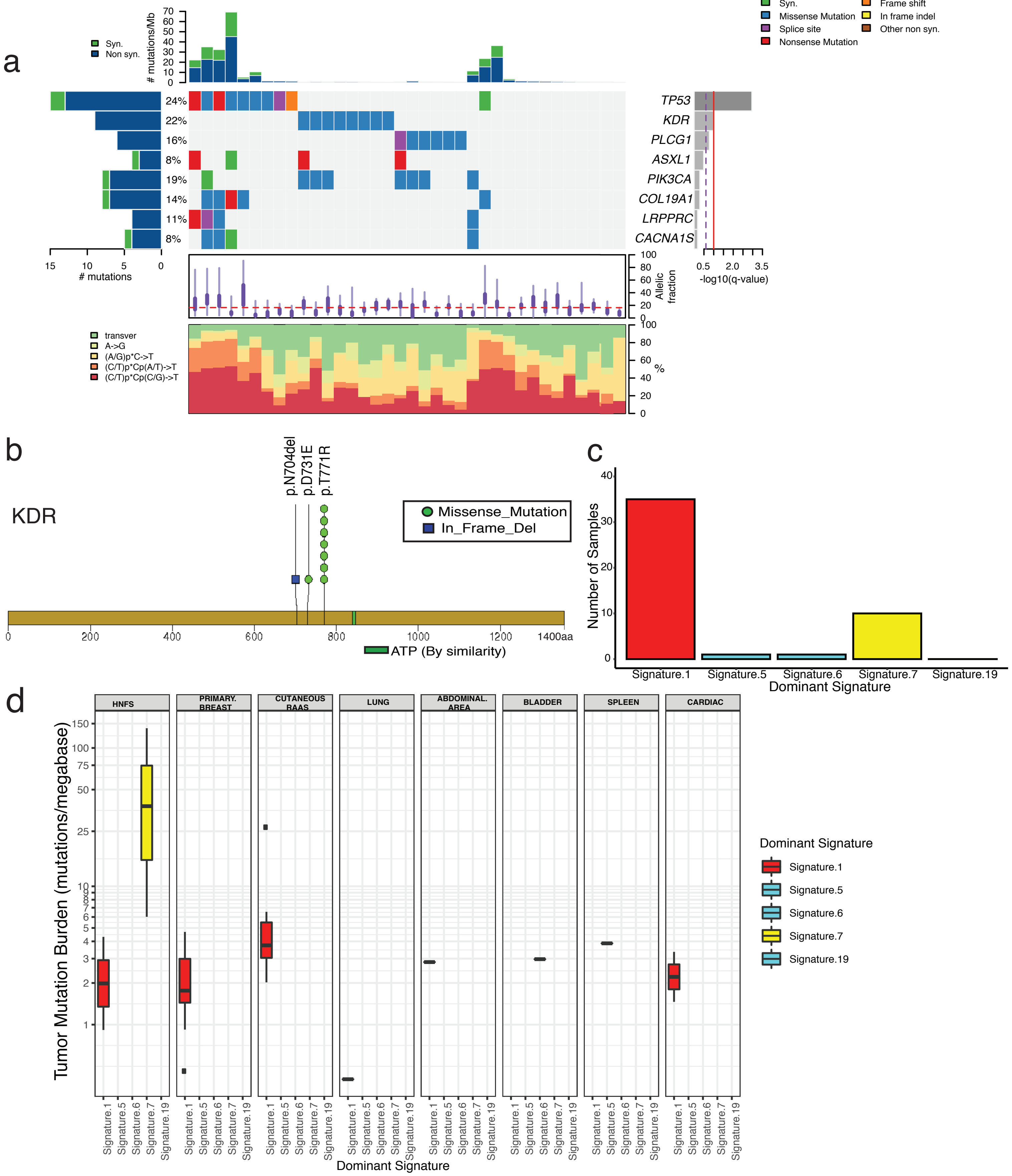


Supplemental Figure 6 - Recurring alterations and dominant mutational signatures in angiosarcoma (a) Co-mutation plot shows significantly recurring mutated genes among 36 AS patients. *TP53* and *KDR* are significantly mutated across the cohort. Genes that have the –log 10 q value >=1 (red line) are significant. (b) Stick plot of KDR showing the recurrent mutations and their positions that were identified in this AS cohort. (c) Bar graph representing the number of tumor samples (y-axis) that are grouped based on 5 dominant mutational signature categories (x-axis). (d) Box plot representing the distribution of TMB (y-axis) across five mutational signature processes (as defined by COSMIC) identified in this cohort and categorized among 8 AS subclassifications (x-axis). Samples with dominant signature 7 (yellow) which corresponds to UV light exposure, have the highest median TMB and this signature was only observed in HNFS AS. AS, Angiosarcoma; TMB, tumor mutation burden, HNFS; head,neck,face,scalp

### Supplemental Figure 7

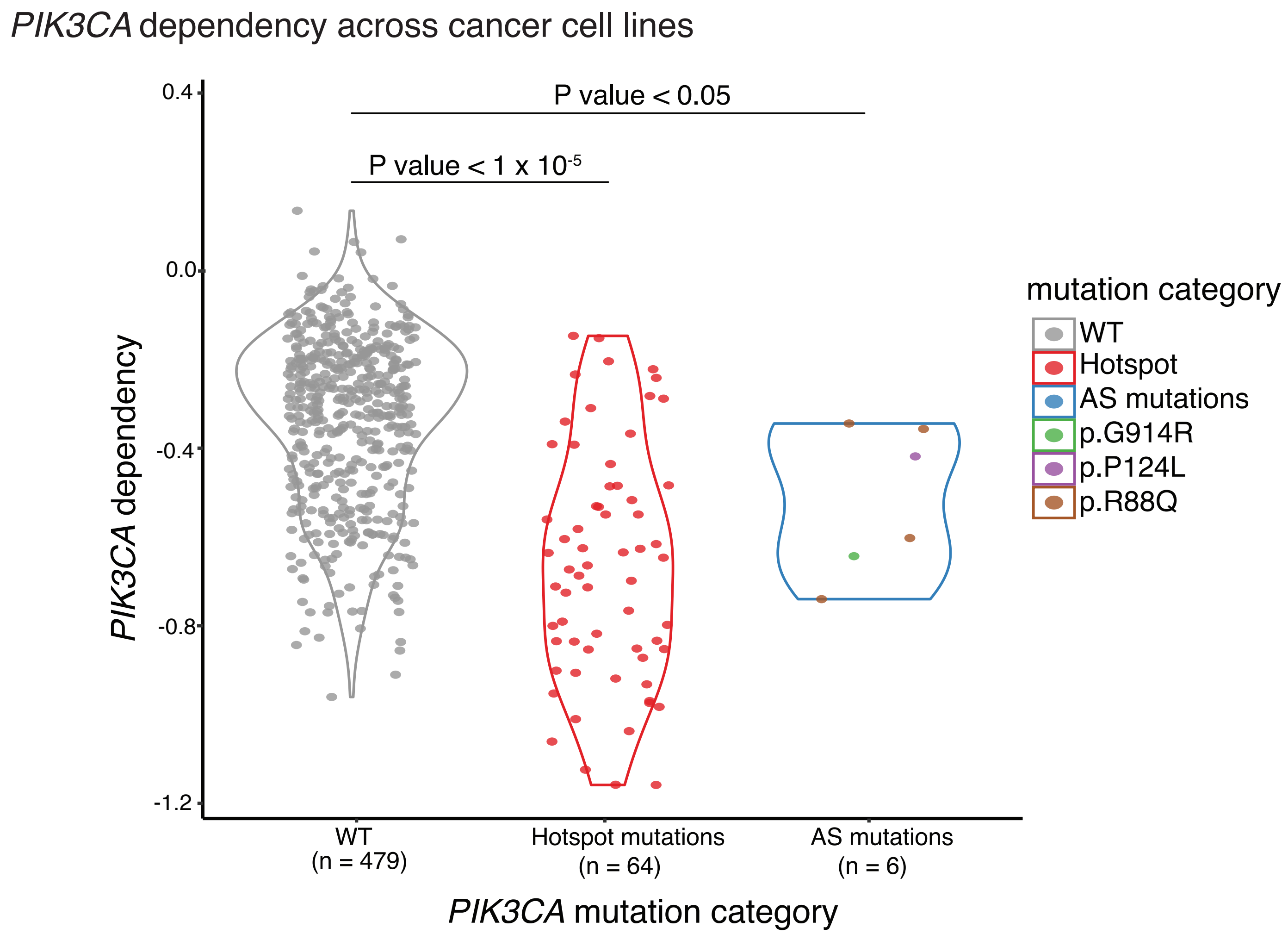


Supplemental FIgure 7. *PIK3CA* mutations found in primary breast angiosarcoma are likely activating. Sensitivity to CRISPR knockout-induced loss of *PIK3CA* was calculated for three groups of cancer cell lines using the Dependency Map dataset (depmap.org): lines containing (1) wild-type *PIK3CA* (gray), (2) *PIK3CA* hotspot mutations (red), and (3) *PIK3CA* mutations seen in AS patients (R88Q [4 lines], P124L [1 line], G914R [1 line]; (colored)). *PIK3CA* hotspot mutations were defined (depmap.org) using their frequency of occurrence in TCGA and COSMIC databases. Relative to cell lines with wild-type *PIK3CA* , *PIK3CA* mutant cell lines are significantly more sensitive to CRISPR knockout-induced loss of *PIK3CA* (*PIK3CA* hotspot mutant lines, p-value< 1X $10^{-5}$; lines with *PIK3CA* with AS mutations, p-value<0.05).
WT, wild-type; AS, angiosarcoma.

Supplemental Figure 8

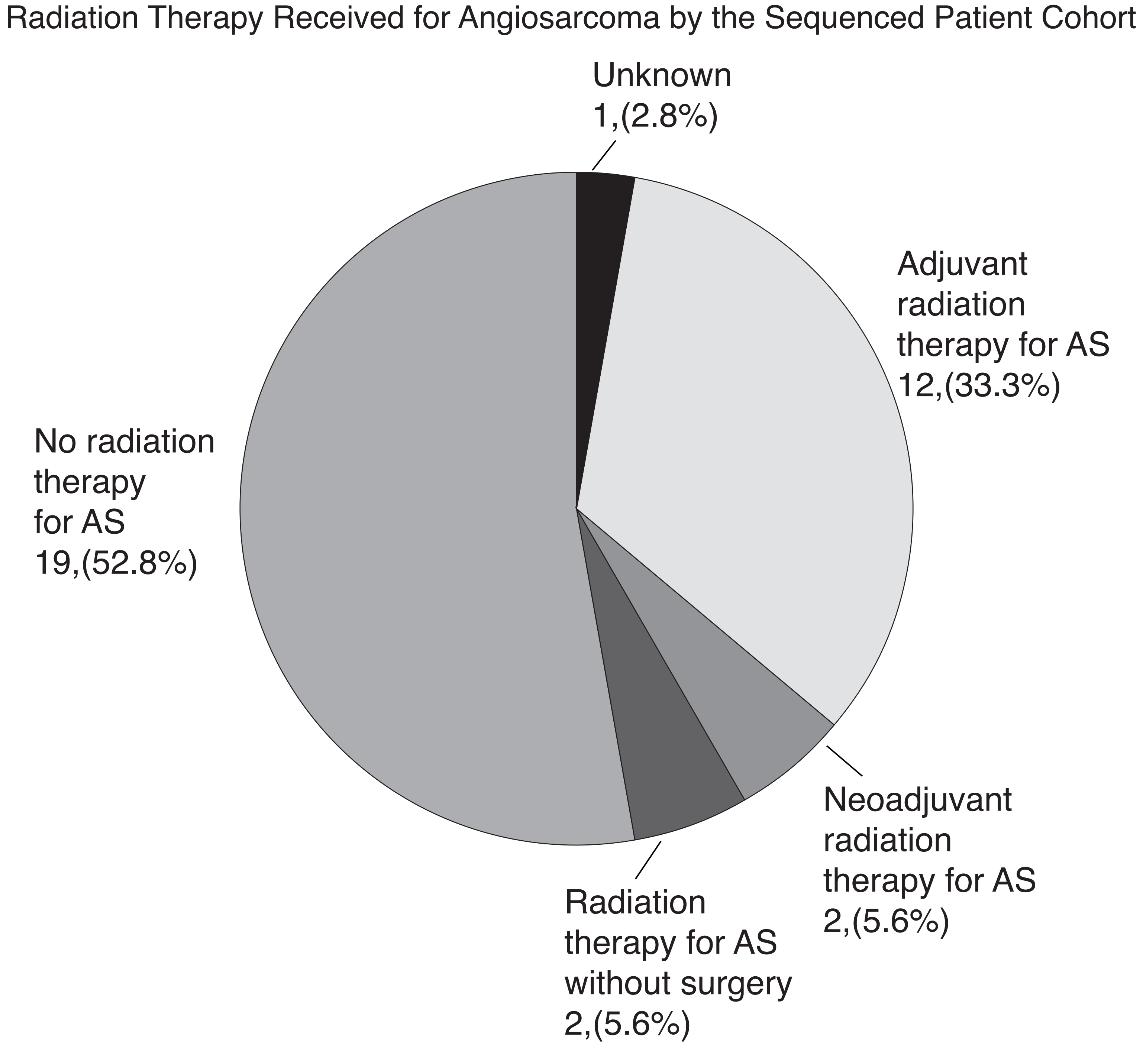


Supplementary Figure 8: Radiation therapy received for angiosarcoma by the sequenced patient cohort. A pie chart showing information regarding radiation treatments received by the 36 sequenced patients in the cohort. This information was abstracted from obtained medical records. These radiation therapies may have been given in conjunction with pharmacological and/or surgical interventions. Patients were categorized into groups depicted in pie chart: patients who received neoadjuvant radiation (2 patients), patients who received adjuvant radiation (12 patients), patients who received radiation for AS without surgery (2), and patients who did not receive radiation for AS (19). For one patient who was categorized as 'Unknown', the medical records were not sufficient to abstract this information.
AS, angiosarcoma.

### Supplemental Figure 9

#### Angiosarcoma Project Patient Referral Source (N=227)

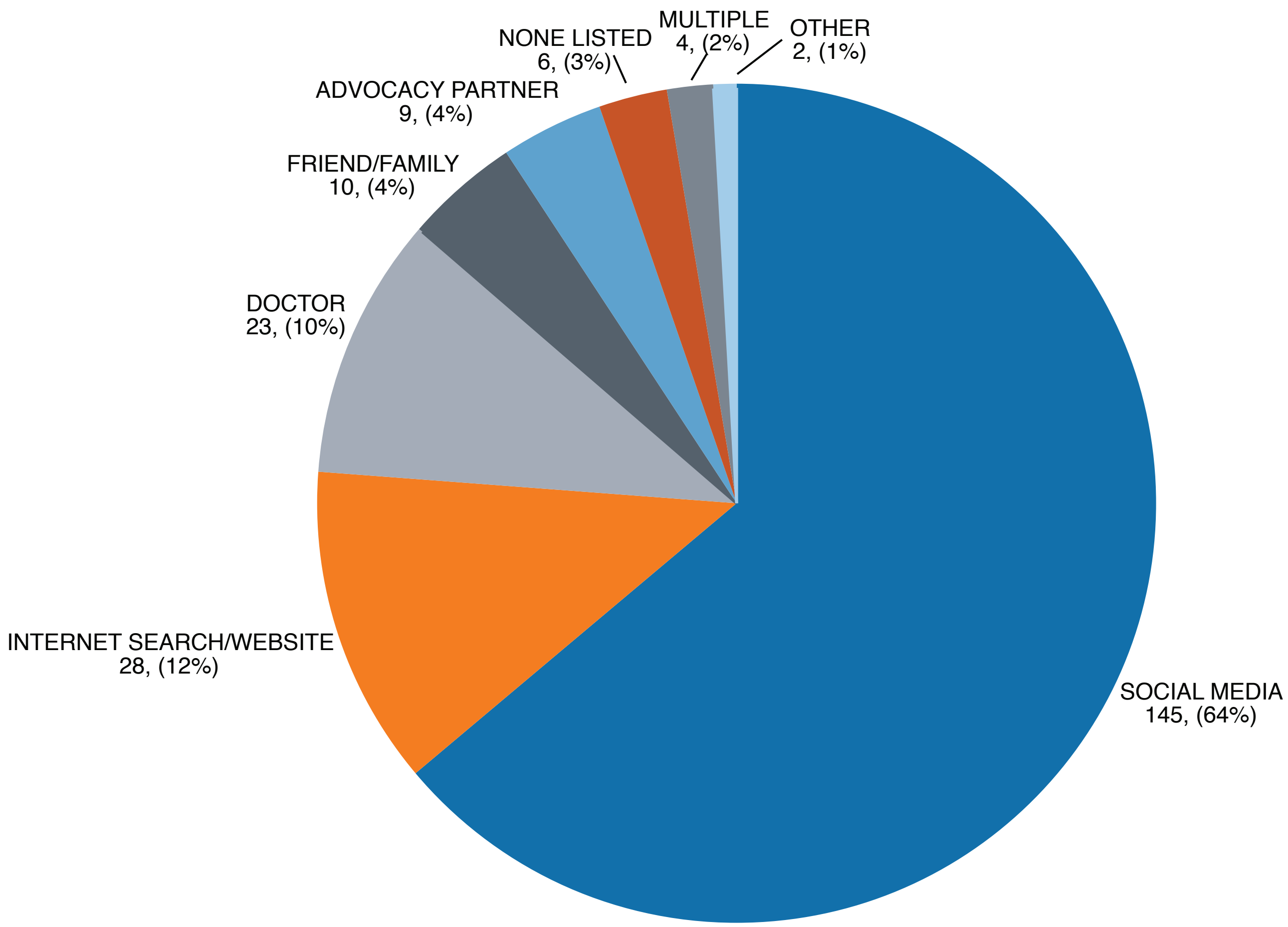


Supplemental Figure 9. Patient referral source to the Angiosarcoma Project. A pie chart showing patient responses to the patient intake survey question 'How did you hear about The Angiosarcoma Project?.' Patient intake surveys completed by the 227 patients who consented for the Angiosarcoma Project as of September 30, 2018 were analyzed. Each free text patient response was grouped into the categories depicted in pie chart. Unique responses from 2 patients were grouped together as 'Other.' 4 patients reported more than one referral source ('Multiple'). 6 patients did not respond to the question ('None Listed').
